# SARS-CoV-2 hijacks p38β/MAPK11 to promote virus replication

**DOI:** 10.1101/2021.08.20.457146

**Authors:** Christina A. Higgins, Benjamin E. Nilsson-Payant, Andrew P. Kurland, Chengjin Ye, Tomer Yaron, Jared L. Johnson, Boris Bonaventure, Prithy Adhikary, Ilona Golynker, Maryline Panis, Oded Danziger, Brad R. Rosenberg, Lewis C. Cantley, Luis Martinez-Sobrido, Benjamin R. tenOever, Jeffrey R. Johnson

## Abstract

SARS-CoV-2, the causative agent of the COVID-19 pandemic, drastically modifies infected cells in an effort to optimize virus replication. Included is the activation of the host p38 mitogen-activated protein kinase (MAPK) pathway, which plays a major role in inflammation and is a central driver of COVID-19 clinical presentations. Inhibition of p38/MAPK activity in SARS-CoV-2-infected cells reduces both cytokine production and viral replication. Here, we combined genetic screening with quantitative phosphoproteomics to better understand interactions between the p38/MAPK pathway and SARS-CoV-2. We found that several components of the p38/MAPK pathway impacted SARS-CoV-2 replication and that p38β is a critical host factor for virus replication, and it prevents activation of the type-I interferon pathway. Quantitative phosphoproteomics uncovered several SARS-CoV-2 nucleocapsid phosphorylation sites near the N-terminus that were sensitive to p38 inhibition. Similar to p38β depletion, mutation of these nucleocapsid residues was associated with reduced virus replication and increased activation of type-I interferon signaling. Taken together, this study reveals a unique proviral function for p38β that is not shared with p38α and supports exploring p38β inhibitor development as a strategy towards developing a new class of COVID-19 therapies.

**Importance:** SARS-CoV-2 is the causative agent of the COVID-19 pandemic that has claimed millions of lives since its emergence in 2019. SARS-CoV-2 infection of human cells requires the activity of several cellular pathways for successful replication. One such pathway, the p38 mitogen-activated protein kinase (MAPK) pathway, is required for virus replication and disease pathogenesis. Here, we applied systems biology approaches to understand how MAPK pathways benefit SARS-CoV-2 replication to inform the development of novel COVID-19 drug therapies.

## Introduction

Severe acute respiratory syndrome coronavirus 2 (SARS-CoV-2), the causative agent of the coronavirus disease 2019 (COVID-19) pandemic, has killed millions since it emerged in 2019. Severe COVID-19 cases are associated with excessive lung inflammation that can lead to acute respiratory distress syndrome, respiratory failure, multi-organ failure, and death (1, 2). This excessive inflammation is in part driven by an imbalanced immune response; compared to other respiratory virus infections, SARS-CoV-2 infection leads to a delay in type I interferon (IFN-I) induction, and excessive pro-inflammatory cytokine and chemokine production (3). While vaccines are highly effective at preventing severe illness and death, novel SARS-CoV-2 variants are continuously emerging with the ability to partially escape prior immunity. Currently, three COVID-19 therapies are available: remdesivir, molnupiravir, and PAXLOVID (4–9). While effective, these therapies need to be administered early in infection, and as they all target single viral proteins, they are susceptible escape mutants. Host-directed therapies are attractive alternatives to antivirals as they are more likely to have broad antiviral activity and are less susceptible to resistance. One such therapy is dexamethasone, an immunomodulatory drug that combats inflammation and reduces COVID-19 mortality, bolstering the concept that targeting host pathways is a viable treatment strategy (10).

We previously reported that the p38 mitogen-activated protein kinase (p38/MAPK) pathway becomes activated during infection, and that inhibition of p38 reduces both inflammatory cytokine expression and SARS-CoV-2 replication, suggesting that p38 inhibition may target multiple mechanisms related to SARS-CoV-2 pathogenesis (11). While the mechanisms by which the p38/MAPK pathway regulates inflammation are well described, the mechanism(s) by which it promotes SARS-CoV-2 replication is unknown. Furthermore, the p38/MAPK pathway is comprised of four p38 kinase isoforms and many downstream kinases, but it is not known which kinases at which levels of the p38/MAPK cascade impact SARS-CoV-2 replication (12).

Here we combined genetic and chemical perturbations, and quantitative transcriptomics and proteomics to better understand interactions between the p38/MAPK pathway and SARS-CoV-2 in human lung epithelial cells. We identified p38β as an essential host factor for SARS-CoV-2 replication and found that while p38β inhibition did not impact viral mRNA abundance, it reduced the abundance of viral protein. Additionally, depletion of p38β specifically resulted in a significant induction of pro-inflammatory cytokines and IFN-I. We applied an unbiased approach to identify novel, putative p38β substrates in the context of SARS-CoV-2 infection, and identified which substrates affect virus replication through siRNA screening. Lastly, we discovered four phosphosites on SARS-CoV-2 nucleocapsid protein (N) that were sensitive to the SB203580 p38 kinase inhibitor and found that phosphoablative mutation of these residues attenuated SARS-CoV-2 growth.

## Results

### Comparisons across SARS-CoV-2 proteomics studies reveal pathways consistently regulated across species and cell types

To better understand the host response to SARS-CoV-2 infection, we quantified changes in protein and phosphosite abundance in A549 human lung epithelial cells expressing ACE2 (A549-ACE2) infected with SARS-CoV-2 at a multiplicity of infection (MOI) of 0.1 for 24 hours (Figure 1A). In total, this analysis comprised 6,089 unique protein groups and 16,032 unique phosphosite groups (Figure 1B). “Phosphosite group” refers to modified residues identified on peptides with sequences that are unique for a single protein or shared across a group of homologous proteins. Phosphosite groups also separate phosphosites identified on singly, doubly, or triply phosphorylated peptides. Throughout this study, we considered protein/phosphosite groups with |log_2_fold-change| > 1 and p-value < 0.05 to be differentially abundant. At the protein level, SARS-CoV-2 proteins (N, ORF1AB/NSP1, ORF1AB/NSP3, ORF9B, and S) were increased by these criteria, with limited changes in host protein groups, consistent with SARS-CoV-2-mediated suppression of host protein synthesis (13). We observed more changes in the phosphoproteome, with 98 and 33 phosphosite groups increased and decreased, respectively (Figure 1C, Table S1).

**Figure 1:**
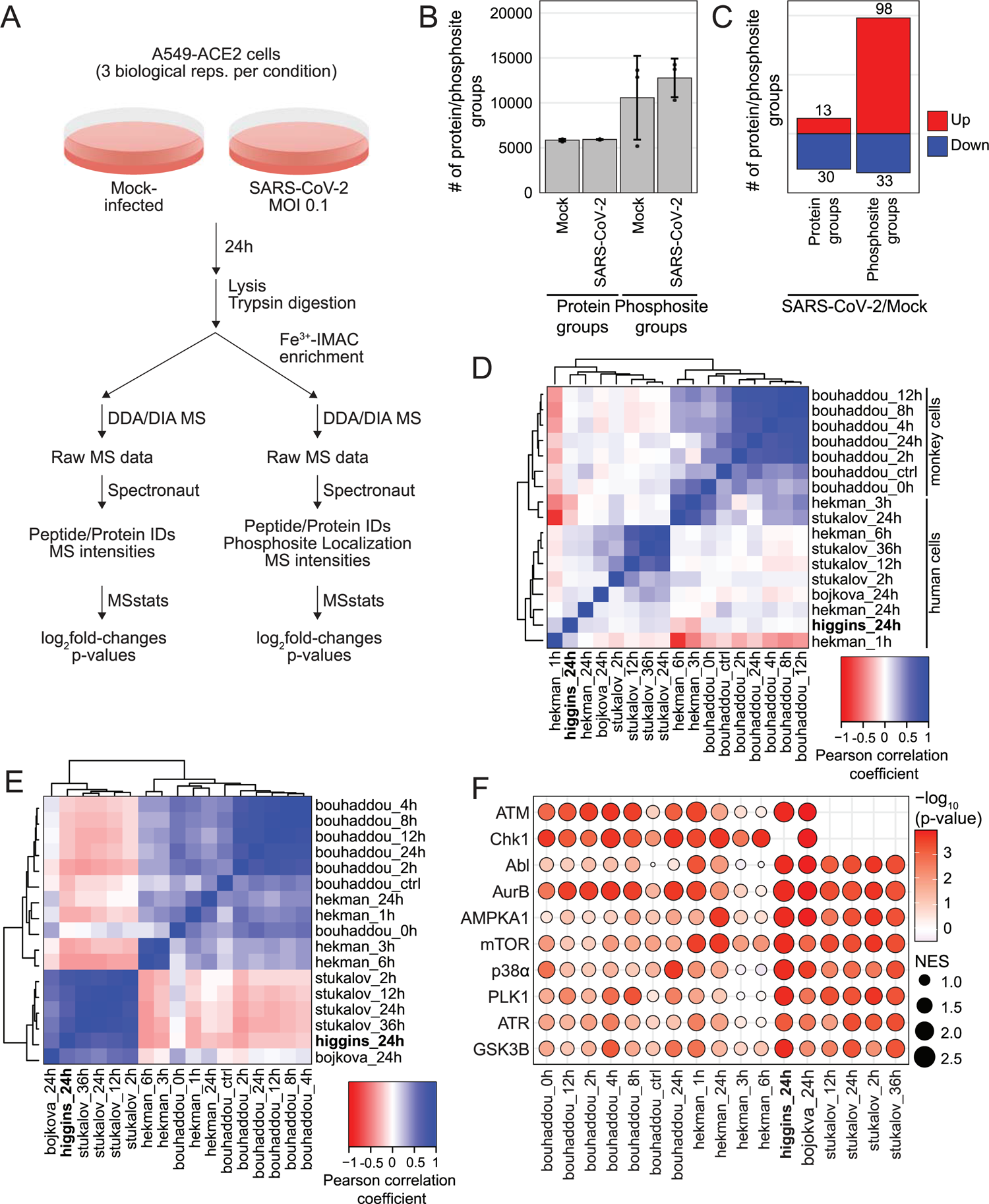
Comparisons across SARS-CoV-2 proteomics studies reveal pathways consistently regulated across species and cell types. A) Schematic of experimental design; B) Plot of the average number of protein groups or phosphosite groups quantified in each condition; error bars represent one standard deviation from the mean for three biological replicates; C) Plot of the number of differentially expressed protein groups or phosphosite groups for SARS-CoV-2-infected cells fold over mock-infected cells; significant change in abundance of protein group or phosphosite group defined as |log_2_(fold-change)| > 1 and p-value < 0.05; D) Heatmap of pairwise Pearson coefficients for phosphosite group log_2_(fold-change) profiles from this study and published studies indicated; E) Heatmap of pairwise Pearson coefficients based on log_10_(p-value) from kinase activity analysis based on log_2_(fold-change) profiles from this study and published studies indicated, also see Table S2; F) Bubble plot of kinase activity analysis based on phosphosite group log_2_(fold-change) profiles for top 10 regulated kinases from this study and published studies indicated; the absolute value of the normalized enrichment score (NES) is indicated by node sizes and the -log_10_(p-value) is indicated by the color scale.

We next compared our A549 data (“higgins”) with four published proteomics studies of SARS-CoV-2 infection of the following cell types: Caco-2 human lung epithelial cells (“bojkova”), Vero E6 African Green Monkey kidney cells (“bouhaddou”), human induced pluripotent stem cell-derived alveolar epithelial type 2 cells (iAT2, “hekman”), and A549 cells (“stukalov”) (11, 14–16). Pairwise Pearson correlation analysis of log_2_fold-change profiles for both protein abundance (Figure S1A) and phosphorylation (Figure 1D) data clustered primarily according to the animal species. Comparing data collected 24-hours post-infection in each study, four phosphosites were upregulated by at least 2-fold in four of the studies: HSPB1/HSP27 S15, MATR3 S188, TRIM28/TIF1B/KAP1 S473, and SZRD1 S107 (Table S2). None of these sites were detected in iAT2 cells. HSPB1 S15 and TRIM28 S473 are canonical p38/MAPK pathway substrates (17–19).

We next performed kinase activity analysis based on log_2_fold-change profiles using a gene set enrichment analysis (GSEA) approach with kinase-substrate annotations from PhosphoSite Plus (Table S3) (20–22). Human cell line kinase activity profiles were strongly correlated (Figure 1E). The ten most regulated kinases across all datasets examined were AURKB, mTOR, CHK1, PLK1, GSK3β, p38ɑ, AMPKA1, ATM, ATR, and Abl (Figure 1F). Kinases involved in cell cycle arrest, ATM, ATR, PLK1, and AURKB, were regulated in all datasets, consistent with evidence that SARS-CoV-2 infection leads to cell cycle arrest (11). Of particular interest to this study, the p38ɑ MAPK was significantly regulated in all studies (Table S3).

### Multiple p38/MAPK pathway components impact SARS-CoV-2 infection in human lung epithelial cells

A gap remains in understanding which components of the p38/MAPK pathway impact SARS-CoV-2 replication. To address this, we employed siRNA screening methodology to assess how depletion of individual p38/MAPK pathway kinases affects SARS-CoV-2 replication. A549-ACE2 cells were transfected with siRNA pools targeting kinase genes, non-targeting control (NTC), or SARS-CoV-2 nucleocapsid (N), and infected with 0.1 MOI of SARS-CoV-2 for 36 hours. The cells were then fixed and stained for SARS-CoV-2 N protein, and the percentage of N-positive cells was determined by immunofluorescence image cytometry, and then normalized to the control infected condition (siNTC-transfected and SARS-CoV-2 infected; Figure 2A). We first screened the four p38 isoforms (p38ɑ/MAPK14, p38β/MAPK11, p38γ/MAPK12, and p38δ/MAPK13) as functional differences between the isoforms in the context of virus infection are particularly understudied. We identified p38β as an essential host factor for SARS-CoV-2 infection, with an approximate 90% reduction in infection and 1000-fold reduction in virus titer when p38β was depleted compared to the infected control condition. Surprisingly, knockdown of p38ɑ did not affect infection even though p38ɑ and p38β are often presumed to be functionally redundant and p38ɑ is thought to be the major isoform regulating immune responses (23). We also found that p38δ depletion reduced infection by approximately 40% and that p38γ knockdown increased it by about 50% (Figure 2B). Based on mRNA-sequencing analysis of A549-ACE2 cells, p38ɑ is the most abundant isoform transcript, followed by p38γ, p38β, and lastly, p38δ (Figure S2A). We then screened the kinases canonically downstream, MAPKAPK2/MK2, MAPKAPK3/MK3, MAPKAPK5/MK5, MSK1, MSK2, and MKNK1, to test the hypothesis that downstream kinase(s) mediate the proviral activity of p38β. MSK2 knockdown reduced infection by approximately 65% and virus titer by 50-fold, while depletion of the other downstream kinases had no effect (Figure 2C-D). We confirmed efficient siRNA knockdown of p38ɑ, p38β, p38γ, and MSK2 protein expression by western blotting (Figure 2E), but were unable to verify knockdown of p38δ with commercial antibodies, likely because its basal expression is low in A549 cells. Cell viability after siRNA transfection was decreased after p38β, p38δ, and MSK2 knockdown by 10-20%, but it is unlikely that infection phenotypes observed are solely due to the decrease in cell viability because other targets like MAPKAP3 also decreased cell viability but did not affect infection (Figure S2B). To validate that the proviral p38β phenotype was not an off-target effect of the siRNAs, we replicated our findings with an independent set of controls and p38β gene-targeting pooled siRNAs (Figure S2C-D).

**Figure 2:**
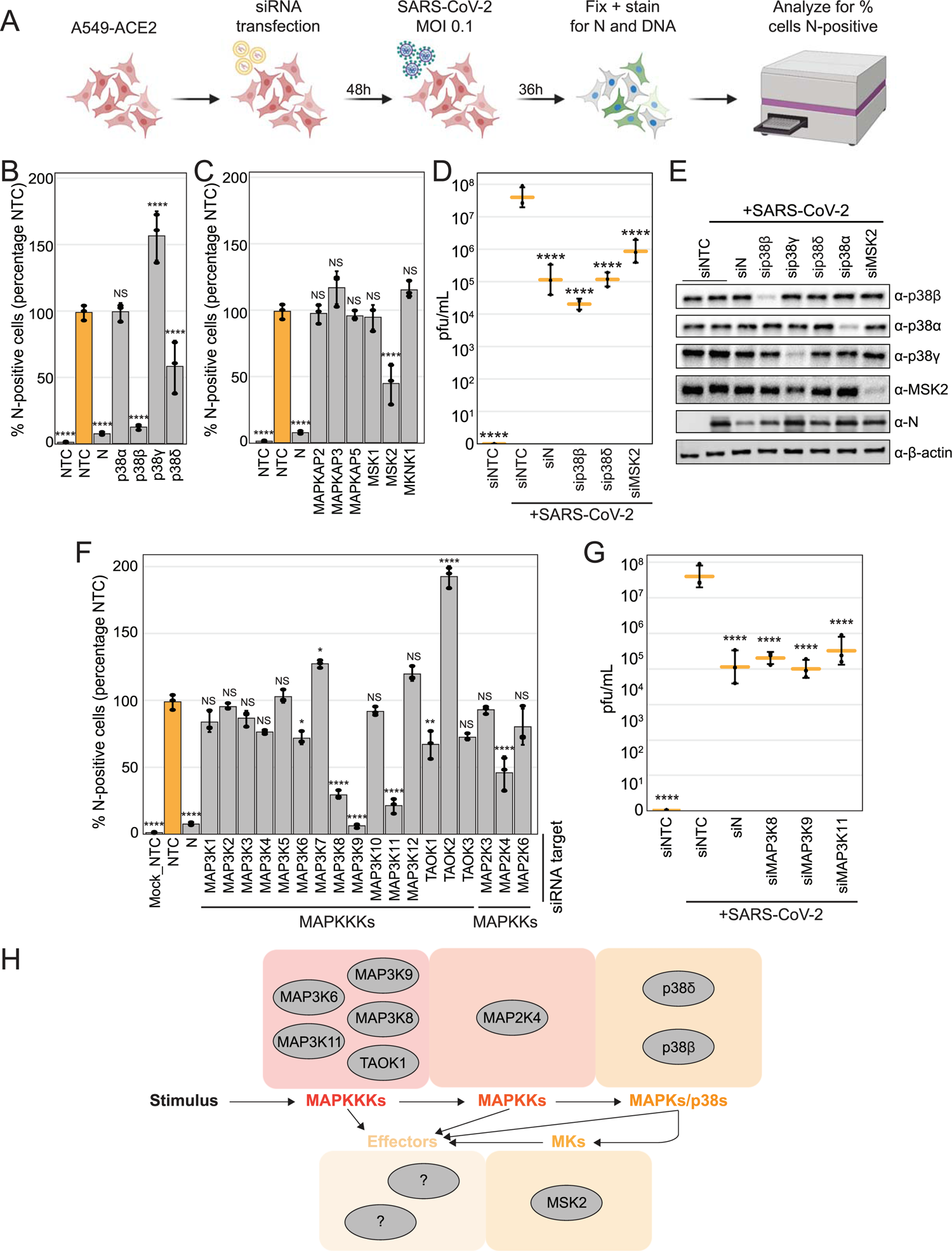
Multiple components of the p38/MAPK pathway impact SARS-CoV-2 infection in human lung epithelial cells. A) Schematic of experiment workflow; B) Plot of the percent of SARS-CoV-2 N-positive cells analyzed using immunofluorescence cytometry, represented as a percentage compared to the siNTC-transfected/SARS-CoV-2-infected control condition, for each indicated p38 isoform or downstream kinase siRNA transfection after SARS-CoV-2 infection at an MOI of 0.1 for 36h in A549-ACE2 cells; C) Plot of SARS-CoV-2 plaque-forming units (pfl)/mL in the supernatant collected from cells in 2B; D) Western blot of cell lysates collected in parallel with cells from 2B; E) Plot of the percent of SARS-CoV-2 N-positive cells, represented as a percentage compared to the siNTC-transfected/SARS-CoV-2-infected control condition, after SARS-CoV-2 infection at an MOI of 0.1 for 36h in A549-ACE2 for each indicated MAPKKK or MAPKK siRNA transfection; F) Plot of SARS-CoV-2 pfu/mL in the supernatant collected from cells in 2E; G) Schematic of p38/MAPK pathway signal transduction highlighting proviral hits from 2B and 2E screens; black arrows indicate a possible phosphorylation event; all error bars represent one standard deviation from the mean for three biological replicates; all p-value annotations were calculated using a one-way ANOVA test with post hoc testing using Tukey’s method comparing each condition to the control infected condition for three biological replicates; “****” = p-value < 0.0001, “***” = 0.0001 < p-value < 0.001, “**” = 0.001 < p-value < 0.01, “*” = 0.01 < p-value < 0.05, “NS” = p-value > 0.05

Focusing next on upstream portions of the p38/MAPK pathway, we tested individual MAPKKK or MAPKK knockdown on virus replication. Among the MAPKKKs screened, we found that MAP3K6/ASK2, MAP3K8/TPL2/COT, MAP3K9/MLK1, MAP3K11/MLK3, and TAOK1/MAP3K16 depletion reduced the percentage of SARS-CoV-2-infected cells by 30-95%, while MAP3K7/TAK1 and TAOK2/MAP3K17/PSK knockdown increased it by nearly 30% and 100%, respectively (Figure 2F). As for the MAPKKs, the canonical p38-regulating MAP2K3/MEK3 and MAP2K6/MEK6, had no phenotype, likely because they are functionally redundant and may not exhibit a phenotype when knocked down individually (24). Interestingly, we found that depletion of MAP2K4/MEK4, widely considered a major regulator of JNK/MAPK signaling (24), decreased infection. This finding could suggest that MAP2K4 can regulate the p38s, or alternatively, that JNK-mediated signaling is involved in SARS-CoV-2 replication. Of the upstream hits, cell viability was affected only by MAP3K8 knockdown (Figure S2B). Finally, we confirmed that virus titers were significantly reduced upon MAP3K8, MAP3K9, and MAP3K11 depletion (Figure 2G). In summary, we found that SARS-CoV-2 replication is promoted by the p38/MAPK signaling cascade specifically involving, when tested individually, the MAPKKKs: MAP3K6, MAP3K8, MAP3K9, MAP3K11, and TAOK1; the MAPKK: MAP2K4; the p38s: p38β and p38δ; and the mediator kinase, MSK2.

### p38β knockdown reduces viral protein, not viral mRNA, in human lung epithelial cells, and promotes type 1 interferon activity

To begin characterizing p38β activity during infection, viral subgenomic RNA (sgRNA), genomic RNA (gRNA), and protein abundance were measured after a high MOI, single-cycle SARS-CoV-2 infection in A549-ACE2 cells transfected with siRNAs targeting controls or p38β. A single cycle infection allowed us to observe the behavior of the virus during one life cycle without multiple iterations of infections confounding observations. First, as a control, knockdown of SARS-CoV-2 N compared to NTC resulted in significant decreases in both viral protein and transcript abundance. p38β depletion resulted in no change in genomic RNA (i.e. *NSP14*) or subgenomic mRNA (i.e. *TRS-N*) abundance, however, knockdown of p38β resulted in a significant decrease in viral protein, but no change in viral transcript abundance (Figure 3A). These findings suggest the mechanism in which p38β promotes virus replication is acting after viral RNA synthesis.

**Figure 3:**
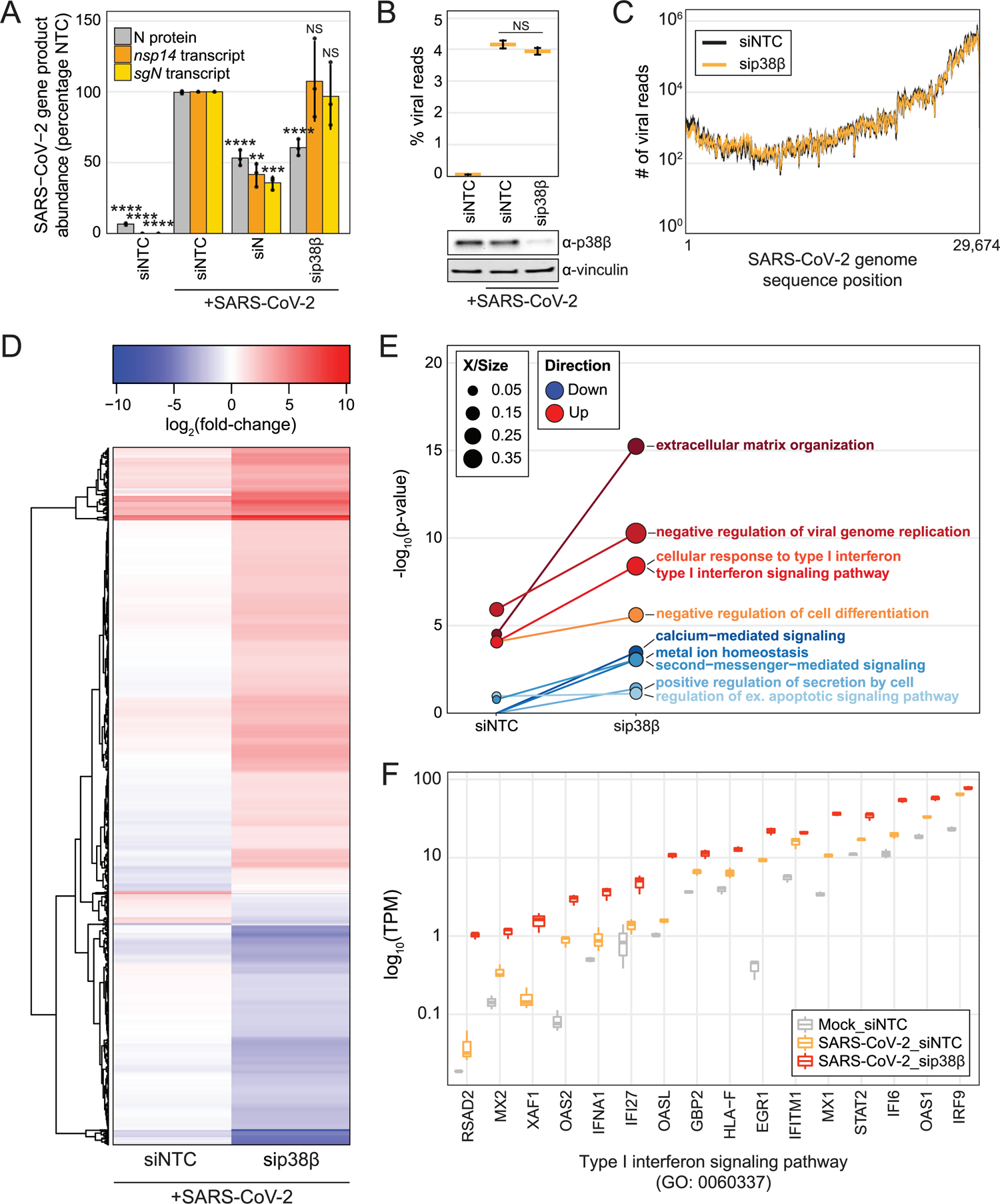
Analysis of cells infected with SARS-CoV-2 for a single virus lifecycle reveals p38β depletion reduces viral protein abundance, but not viral RNA abundance, and promotes type 1 interferon activity. A) Plot of the percentage of SARS-CoV-2 N protein-positive cells analyzed using immunofluorescence microscopy (grey) or *NSP14* or *sgN* transcript abundance detected using RT-qPCR (gold/yellow) normalized to the control infected condition after SARS-CoV-2 infection at an MOI of 2 for 8h in A549-ACE2 cells; error bars represent one standard deviation from the mean; p-values were calculated using a one-way ANOVA test with post hoc testing using Tukey’s method comparing each condition to the control infected condition; “****” = p-value < 0.0001, “***” = 0.0001 < p-value < 0.001, “**” = 0.001 < p-value < 0.01, “*” = 0.01 < p-value < 0.05, “NS” = p-value > 0.05; statistics for protein generated using three biological replicates and statistics for mRNA abundance generated using nine biological replicates; B) Plot of percent viral reads from mRNA-Seq of A549-ACE2 cells transfected with siNTC or sip38β and infected with SARS-CoV-2 MOI 0.75 or mock-infected for 8 hours; significance annotated as p-value from one-way ANOVA test with post hoc testing using Tukey’s method comparing each condition to the control infected condition; NS = p-value > 0.05; western blot from lysates collected in parallel below; C) Hierarchically clustered heatmap of differentially expressed genes for sip38β-transfected/SARS-CoV-2 infected compared to siNTC-transfected/SARS-CoV-2-infected, but shown here as each infected condition fold over siNTC-transfected/mock-infected; rows represent each gene; color corresponds to log_2_(fold-change) as indicated; D) Plot of four most significant GO terms enriched from sip38β-transfected/SARS-CoV-2-infected fold over siNTC-transfected/SARS-CoV-2-infected differentially upregulated (red shades) or downregulated (blue shades) genes; point size represents was proportion of total genes associated with a GO term are represented; E) Plot of log_10_(transcripts per million) for each gene represented in the indicated GO term for each condition, from same analysis as 3D. All raw and processed mRNA-Seq data is available on NCBI GEO GSE183999.

We next used mRNA sequencing to analyze transcriptome changes in p38β-depleted or control A549-ACE2 cells infected for a single replication cycle of SARS-CoV-2. Samples clustered by condition as measured by principal component analysis (S3A-B). Control infected cells fold over mock-infected cells resulted in 197 differentially expressed genes (DEGs, |log_2_fold-change| > 1.5 and adj. p-value < 0.05) and p38β-depleted, infected cells fold over mock resulted in 1,303 DEGs (Figure S3C-D, Table S5). Consistent with the RT-qPCR results, the percentage of viral reads did not change between control infected cells and p38β-depleted cells, even though p38β protein expression was significantly reduced (Figure 3B).

937 genes were differentially expressed between the infected, p38β-depleted cells, and the control infected cells (Table S5). These genes, shown as a heatmap of log_2_fold-changes for each condition fold over mock (Figure 3D), frequently trended in the same direction, but to a larger degree with p38β knockdown. Additionally, gene ontology (GO) enrichment analysis revealed that extracellular matrix organization- and IFN-I-related GO terms were significantly enriched by the upregulated DEGs (Figure 3E). These data suggest that p38β negatively regulates the expression of pro-inflammatory cytokines and IFN-I, which was not expected as p38 kinases are generally thought to positively regulate cytokine expression. The same analysis on the downregulated DEGs enriched for GO terms related to second messenger-mediated signaling, metal ion homeostasis, and cell secretions (Figure 3E, Table S6). Focusing on the genes that contributed to the significance of “Type 1 interferon signaling pathway”, some genes were impacted more than others by p38β depletion compared to infected control and mock-infected conditions. For example, the transcripts per million reads (TPM) counts for genes such as *RSAD2*, *IFNA1, IFI27*, and *OASL* were similar for mock-infected and infected control, but much higher in the p38-depleted condition, whereas for most other genes, both infected conditions had more counts than the mock (Figure 3F). These data suggest p38β may differentially regulate the expression of IFN-related genes. Genes that contributed to the most significantly enriched annotations, extracellular matrix organization and calcium-mediated signaling, exhibited similar abundance changes (S3E-F).

### p38β proviral mechanism is primarily STAT1-independent but leads to ISG expression as a byproduct

To assess if transcriptome changes were also reflected at the proteome level, we assessed how siRNA knockdown of p38β or MSK2 in the context of SARS-CoV-2 infection affects the host proteome using quantitative proteomics. In biological quadruplicate, A549-ACE2 cells were transfected with pooled siRNAs targeting NTC, p38β, or MSK2. Cells were then infected with SARS-CoV-2 and 36-hours post-infection, cells were lysed and subjected to quantitative proteome and phosphoproteome analysis (Figure 4A). A total of 4,900 unique protein groups and 14,414 unique phosphosite groups were identified (Figure 4B and Table S1). These data clustered by their respective siRNA targets in Pearson correlation analysis and each sample had similar normalized log_2_intensity distributions (Figure S4A-D). In comparison to siNTC-transfected/mock-infected cells, siNTC-transfected/SARS-CoV-2-infected (control infected) cells yielded few changes to the proteome and large changes to the phosphoproteome with approximately 1,000 phosphosite groups significantly changing. More changes were observed here than in our preliminary A549-ACE2 analysis (Figure 1); the siRNA analysis included a greater number of biological replicates (4 vs. 3) and reduced technical variability that yielded a greater number of features with lower p-values. Furthermore, siRNA transfection may have impacted the number of significant changes.

**Figure 4:**
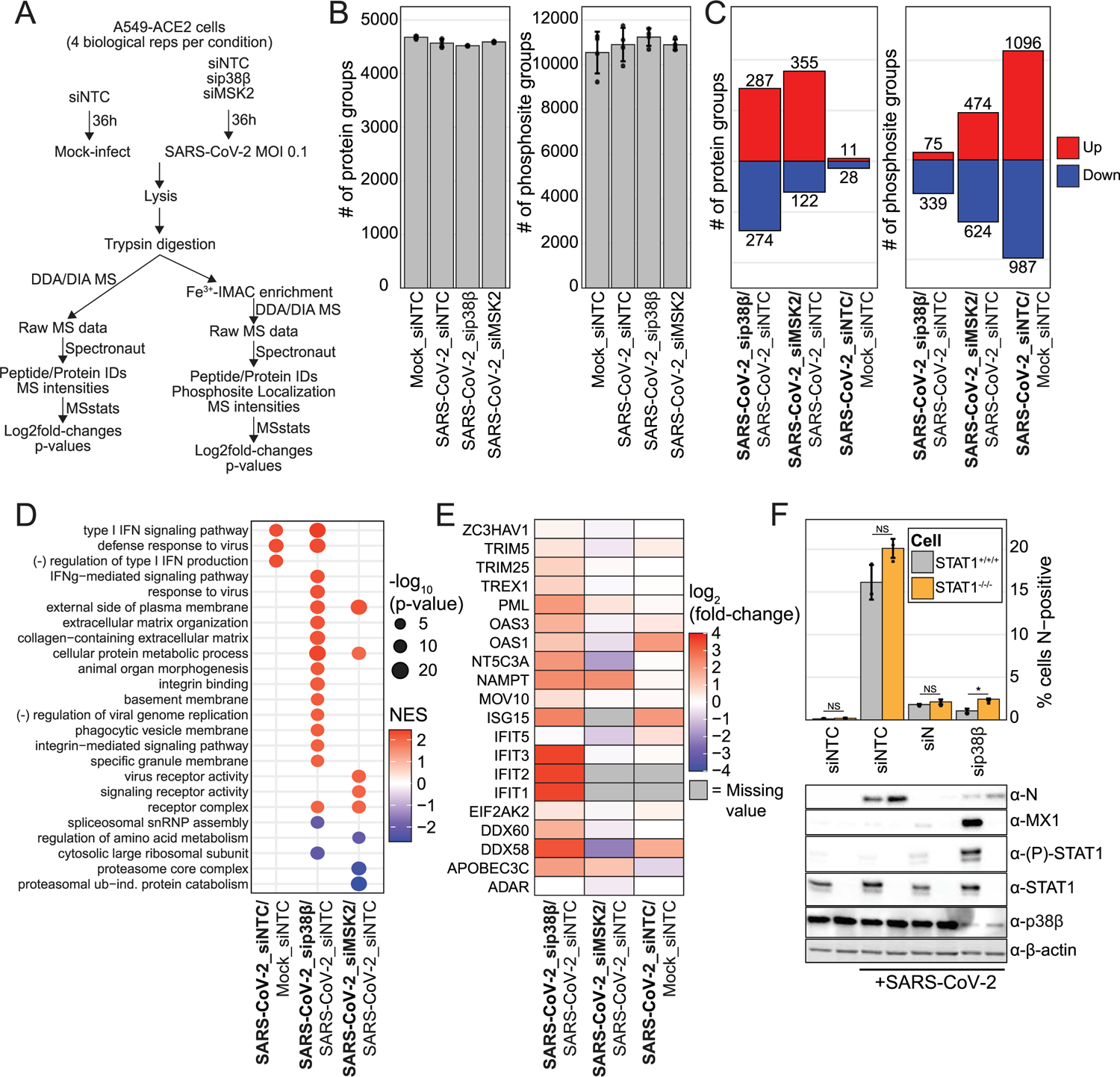
p38β proviral mechanism is primarily STAT1-independent but leads to ISG expression as a byproduct. A) Schematic of experiment workflow; B) Plot of the number of protein groups or number of phosphosite groups identified in each biological replicate; error bars represent one standard deviation from the mean for four biological replicates; C) Plot of the number of significantly differentially abundant protein groups and phosphosite groups for condition comparisons as indicated; significant change in abundance of protein group or phosphosite group defined as |log_2_(fold-change)| > 1 and p-value < 0.05; D) GO terms enriched from analysis of differentially abundant protein groups for each comparison from protein abundance data using gene set enrichment analysis (GSEA); E) Heatmap of log_2_(fold-change) for each indicated ISG in each indicated condition comparison from protein abundance data; F) Plot of the percent of SARS-CoV-2 N-positive cells analyzed using immunofluorescence cytometry for each indicated transfection condition after SARS-CoV-2 infection at an MOI of 0.1 for 30h in A549-ACE2 cells or A549-ACE2ΔSTAT1; error bars represent one standard deviation from the mean for three biological replicates; p-value annotations were calculated using a one-way ANOVA test with post hoc testing using Tukey’s method comparing each condition between each cell type for three biological replicates; “****” = p-value < 0.0001, “***” = 0.0001 < p-value < 0.001, “**” = 0.001 < p-value < 0.01, “*” = 0.01 < p-value < 0.05, “NS” = p-value > 0.05; below are western blots of lysates collected in parallel with cells analyzed in above plot

Knockdown of each kinase in SARS-CoV-2-infected cells led to substantial changes to the proteome. Compared to control infected cells, knockdown of p38β or MSK2, led to 287 and 355 unique protein groups significantly increasing, respectively, and 274 and 122 protein groups significantly decreasing, respectively. There were also significant changes to the phosphoproteome; in comparison to control infected cells, knockdown of p38β or MSK2 led to 75 and 474 unique phosphosite groups significantly increasing, respectively, and 339 and 624 phosphosite groups significantly decreasing, respectively (Figure 4C, Table S1). Consistent with our observations made at the transcriptome level, GO enrichment analysis of protein abundance log_2_fold-change profiles revealed that SARS-CoV-2-infected cells depleted of p38β exhibited a strong IFN-I signature compared to control infected cells. MSK2 knockdown in SARS-CoV-2-infected cells did not lead to a comparable phenotype, suggesting MSK2 does not mediate p38β-related interferon regulation (Figure 4D, Table S7). Focusing on well-characterized ISGs (25), most were enhanced in response to p38β depletion cells compared to control infected cells (Figure 4E). Western blotting confirmed that p38β knockdown led to an increase in MX1, a prototypical ISG, and specifically in the context of infection (Figure S4E).

These findings led us to question whether perturbation of p38β prevents SARS-CoV-2 replication by inducing ISG expression, which in turn suppresses viral replication, or if perturbation of p38β prevents replication in an independent manner that incidentally leads to the expression of ISGs, for example, by exposing a pathogen-associated molecular pattern (PAMP) that is detected by the innate immune sensors. To address this, we performed SARS-CoV-2 infections and siRNA knockdowns of controls or p38β in A549-ACE2 cells or STAT1-knockout A549-ACE2 cells, which are insensitive to IFN-I or IFN-III signaling. If STAT1-dependent expression of ISGs is required to restrict infection in cells with reduced p38β expression, deletion of STAT1 would restore SARS-CoV-2 replication to wild-type levels. We found that while the reduction in infection when p38β is depleted is significant, it is not appreciably rescued by STAT1-knockout, indicating that the mechanism of action of p38β is primarily STAT1-independent (Figure 4F). We confirmed by western blotting efficient knockdown of p38β, and STAT1-knockout, ablation of MX1 expression and STAT1 phosphorylation in the STAT1-knockout A549-ACE2 cells (Figure 4F). We next tested the JAK1/2 inhibitor ruxolitinib, which acts upstream of STAT1 and is broadly effective at preventing IFN (and other cytokine) activity, and again, observed that JAK1/2 inhibition did not rescue the infection defect associated with p38β knockdown (Figure S4F). These findings demonstrate that the enhanced antiviral response that results following p38β knockdown is not the primary mechanism by which SARS-CoV-2 infection is reduced.

### Quantitative, unbiased phosphoproteomics analysis pipeline identifies novel p38β substrates

In order to identify putative p38β substrates that may explain the mechanism by which p38β promotes infection, we created an analysis pipeline to assess proteome and phosphoproteome data from experiments employing three different p38β perturbation strategies: 1) siRNA knockdown of p38β (described previously); 2) titrated treatment of cells with the SB203580 p38ɑ/β inhibitor beginning one-hour before a 24-hour SARS-CoV-2 infection (pre-treatment), and 3) 24-hour SARS-CoV-2 infection with the last four hours being in the presence of SB203580 (terminal treatment) (Figure 5A).

**Figure 5:**
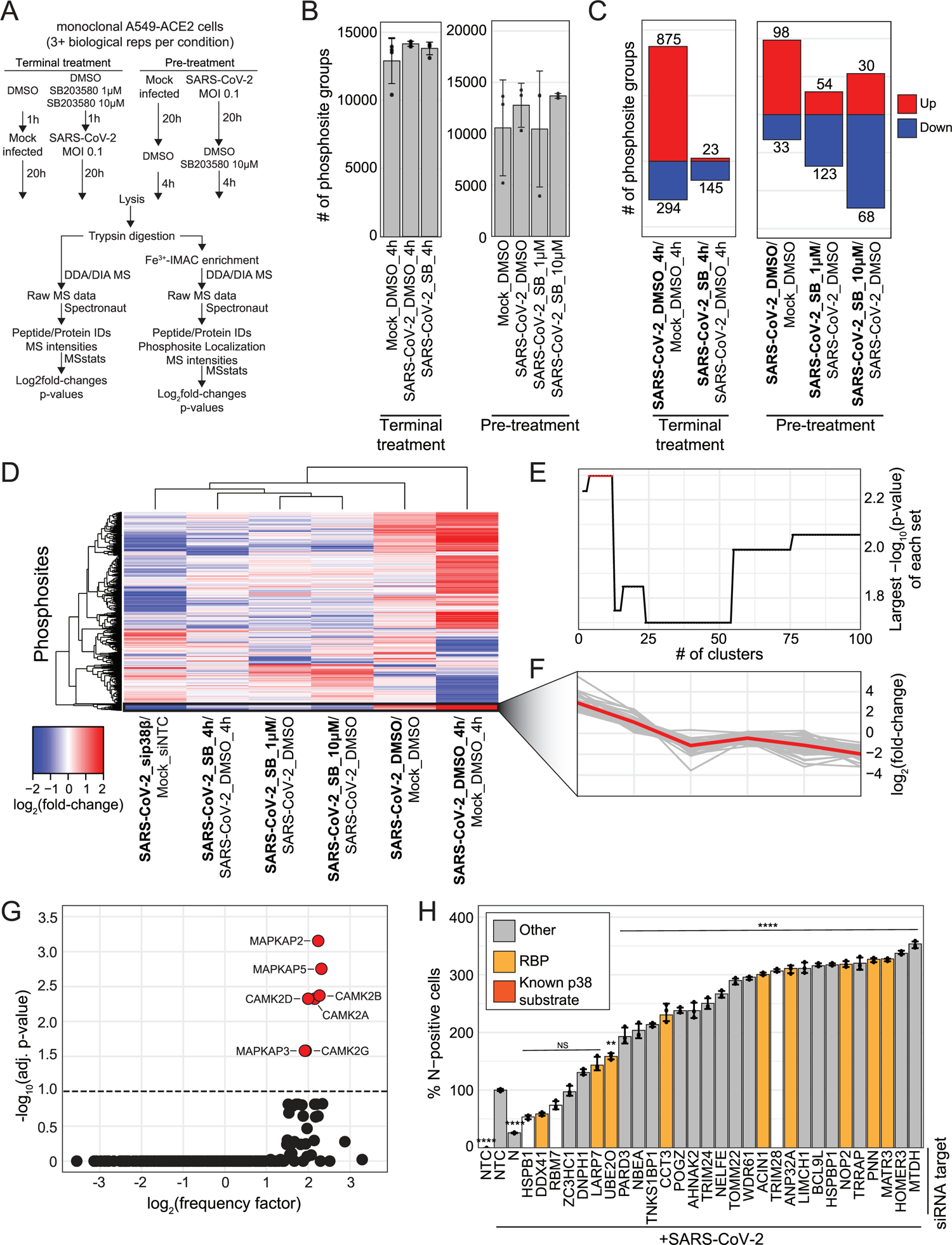
Novel, putative p38 pathway kinase substrates identified using KIPPC analysis pipeline. A) Schematic of experiment workflow; B) Plot of the number of phosphosite groups identified in each condition; error bars represent one standard deviation from the mean for at least three biological replicates; C) Plot of the number of significantly differentially abundant phosphosite groups for each indicated condition comparison for the “terminal treatment” experiment arm (left), or the “pre-treatment” experiment arm (right); significant change in abundance of protein group or phosphosite group defined as |log_2_(fold-change)| > 1 and p-value < 0.05; D) Heatmap of log_2_(fold-change) for differentially abundant phosphosite groups (rows) in each indicated condition comparison (columns), hierarchically clustered; cluster of interest in black box; E) Plot of -log_10_(p-value) of each highest - log_10_(p-value) cluster from each clustering iteration; F) Plot of log_2_(fold-change) profiles of cluster-of-interest phosphosite groups (grey line) for each condition comparison in 5D; red line indicates average profile for all cluster-of-interest phosphosite groups; G) Plot of -log_10_(adjusted p-value) and -log_2_(frequency factor) based on the comparison of cluster-of-interest phosphosite motif sequences to the consensus substrate motif sequence for each characterized human kinase; red points are kinases with significantly similar consensus substrate motifs to the cluster-of-interest phosphosite group sequences, and black points represent kinases with insignificantly similar consensus substrate motifs; H) Plot of percent of SARS-CoV-2 N-positive cells analyzed using immunofluorescence cytometry normalized to the control infected condition for each indicated transfection condition after SARS-CoV-2 infection at an MOI of 0.1 for 30h in A549-ACE2 cells; error bars represent one standard deviation from the mean for three biological replicates; p-values were calculated using a one-way ANOVA test with post hoc testing using Tukey’s method comparing each condition to the control infected condition for three biological replicates; “****” = p-value < 0.0001, “***” = 0.0001 < p-value < 0.001, “**” = 0.001 < p-value < 0.01, “*” = 0.01 < p-value < 0.05, “NS” = p-value > 0.05.

For chemical perturbation strategies, selected SB203580, a well-characterized p38ɑ/β-specific small molecule inhibitor of enzymatic activity as p38β-selective inhibitors are not currently available (26). However, SB203580 is estimated to be ten times more potent at inhibiting p38ɑ as p38β (27). These data cluster by their respective conditions in Pearson correlation analysis and each sample has similar normalized log_2_intensity distributions within each experiment (Figure S4A-H). The terminal drug treatment experiment yielded 16,220 unique phosphosite groups, and the pre-treatment experiment, 16,032 (Figure 5B and Table S1). Pre-treated samples had substantially more changes in the proteome and phosphoproteome compared to terminally treated samples, likely reflecting how the limited drug-exposure time does not allow for significant changes protein expression, and SB203580 pre-treatment reduces infection (Figure 1C, Table S1, 11). We observed a significant decrease in phosphorylation of known p38 substrates: CP131 S47, HSPB1 S15, PARN S557, RIPK1 S320, in response to SB203580 in both pre-treatment and terminal treatment experiments (Figure S5I-J). In the pre-treatment protein abundance data, we did not see an upregulation of ISGs, contrasting with observations made upon genetic perturbation of p38β (Table S1). It is likely that the ISG phenotype did not develop because SB203580 is primarily a p38ɑ inhibitor and SB203580-mediated p38β inhibition was not as effective as genetic inhibition.

Proceeding with our analysis pipeline, we next combined the phosphoproteome profiles with of both drug treatment datasets with the p38β knockdown profiles and developed a supervised hierarchical clustering approach called <u>ki</u>nase <u>p</u>erturbation <u>p</u>hospho-profile <u>c</u>lustering (KiPPC) (Figure S5L). Data were first filtered for single phosphorylated phosphosite groups, no missing values across comparisons, and significant changes in at least one comparison yielding 1,191 total phosphosite profiles, including 12 phosphosites annotated in Phosphosite Plus as substrates of p38α, p38β, or one of their downstream effector kinases (i.e., MK2, MK3, MK5, MSK1, MSK2, or MKNK1) (21). The profiles were then hierarchically clustered based on their Euclidean distances (Figure 5D), generating a dendrogram tree that was then cut iteratively 99 times in order of decreasing height to generate between 2 and 100 clusters. For each iteration, the significance of enrichment of p38α/β substrates in each cluster was calculated with a hypergeometric test. The cluster most significantly enriched for known p38α/β substrates occurred in iterations 3-12, with a hypergeometric p-value of 0.005 (Figure 5E). The phosphosite profiles within this cluster are very similar, with the representative profile behaving as expected: during SARS-CoV-2 infection, the log_2_fold-change is high because the p38 pathway is active, and when p38β is genetically or chemically inhibited, it decreases (Figure 5F). The cluster contains 35 phosphosites in total including three annotated p38α/β substrate sites (HSPB1 S15, RBM7 S137, and TRIM28 S473), several p38α/β substrates at phosphosites not previously annotated as p38α/β-dependent (TRIM28 S471, HSPB1 S78, HSPB1 S82, NELFE S51), as well as proteins physically associated with annotated p38α/β substrates (LARP7 and TRIM24, (Table S8). Additionally, the cluster is enriched for processes commonly associated with the p38/MAPK pathway including RNA binding, protein folding, and transcription elongation (Figure S5M) (12). In support of our KiPPC analysis results, we used the Kinase Library to analyze our cluster’s phosphosite motifs and found that the kinases most likely to phosphorylate these substrates are p38/MAPK pathway members, MAPKAP2, MAPKAP3, and MAPKAP5, and related CAMK-type kinases, CAMK2-A, -B, -D, and -G (Figure 5H) (28). We next aimed to determine if any of the novel, putative substrates impact SARS-CoV-2 replication by employing the same siRNA screening methodology described above. We found that depletion of a vast majority of putative p38α/β substrates tested (22 of 29) resulted in a significant increase in infection (Figure 5G). We also assessed cell viability in response to each siRNA transfection (Figure S5N). While these data do not specifically reflect the impact of phosphorylation at these sites on virus replication, this screen revealed that a large number of putative p38α/β substrates play critical roles in SARS-CoV-2 infection.

### SARS-CoV-2 N protein phosphorylation is sensitive to p38 inhibition

In addition to identifying novel host p38 substrates, we also explored possible p38-dependent phosphorylation of SARS-CoV-2 proteins. In order to focus on viral phosphosites, we specifically looked at the short, terminal drug treatment dataset because this experimental framework does not affect total viral protein abundance, allowing us to directly and accurately quantify changes to viral phosphosites in response to p38 inhibition. In our analyses, SARS-CoV-2 N was the only viral protein identified that harbored SB203580-sensitive phosphosites based on p-values, although the fold-changes were less than the 2-fold threshold we implemented throughout this study. We identified four N phosphosites (S21, S23, T24, and S26) that decreased during infection in response to SB203580 compared to DMSO treatment (Figure 6A and S6A). These sites are located in an intrinsically disordered region close to the N-terminus of N. Additionally, these amino acid residues have no or relatively low entropy (i.e. variation) amongst SARS-CoV-2 variants in the Nextstrain resource (Figure 6A) (29). We confirmed no significant difference in total N protein abundance between DMSO-treated, infected cells and SB203580-treated, infected cells (Figure 6B). These sites, phosphorylated either directly or indirectly by p38ɑ and/or p38β, could confer changes in N activity that affect virus replication. To test this hypothesis, we generated a recombinant, phosphoablative mutant of SARS-CoV-2 USA-WA1/2020 (rSARS-CoV-2^N4A^) containing alanine substitutions at the four SB203580-sensitive phosphoresidues (Figure 6C). In a longitudinal experiment comparing virus titer of recombinant wild-type SARS-CoV-2 USA-WA1/2020 (rSARS-CoV-2^WT^) to rSARS-CoV-2^N4A^, the mutant virus was significantly attenuated in titer at several timepoints (Figure 6D). Interestingly, we also observed that rSARS-CoV-2^N4A^ infection of Vero E6 produced morphologically different plaques, larger with less defined edges, than rSARS-CoV-2^WT^ (Figure 6E). Lastly, rSARS-CoV-2^N4A^ infection of A549-ACE2 cells led to a higher induction of a canonical ISG, MX1, than infection with rSARS-CoV-2^WT^ (Figure 6F). While future work is required to assess the underlying biochemistry responsible for the N4A phenotype, these data clearly demonstrate a need for p38β in SARS-CoV-2 biology. In summary, we found the N-terminal phosphorylation of N promoted viral production, albeit less significantly than p38β knockdown, suggesting that p38 impacts viral replication by modulating both viral and host substrates.

**Figure 6:**
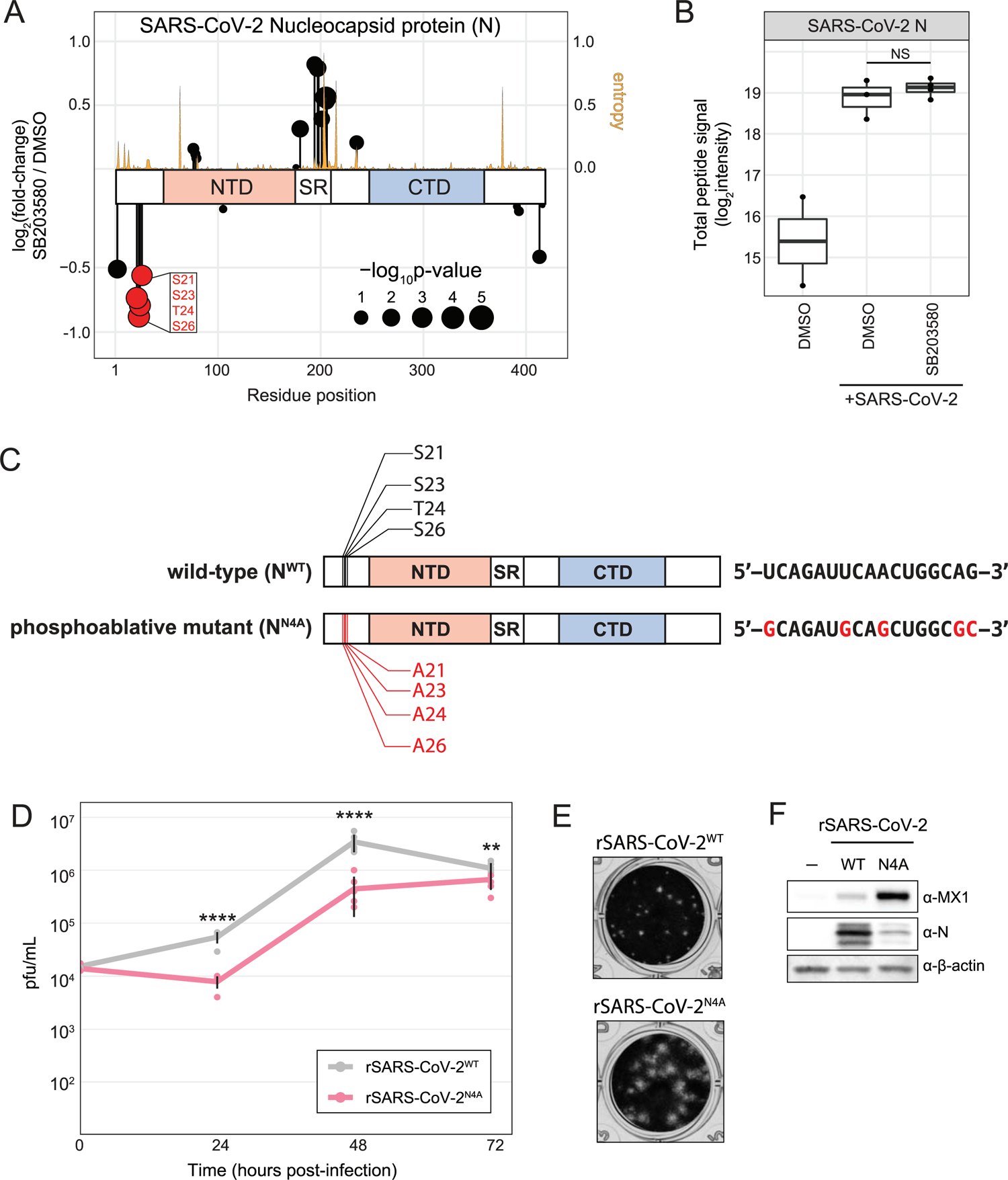
Phosphoablative mutation of four SB203580-sensitive residues on SARS-CoV-2 nucleocapsid protein attenuates virus growth. A) Plot of log_2_(fold-changes) for each phosphosite group on SARS-CoV-2 N differentially abundant for 10μM SB203580-terminally-treated/SARS-CoV-2-infected fold over control infected condition; entropy (amino acid sequence conservation between SARS-CoV-2 variants; higher entropy = less conserved) indicated as yellow line; B) Plot of log_2_(signal intensity) of total N protein abundance from terminal-treatment experiment (Figure 5A); NS = p-value > 0.05; C) Schematic comparing N^WT^ to N^N4A^ (left), and the corresponding nucleotide sequence conferring the mutation (right); D) Graph comparing rSARS-CoV-2^WT^ and rSARS-CoV-2^N4A^ titer (pfu/mL) at several points during an infection performed at an MOI of 0.01 in A549-ACE2 cells; error bars represent one standard deviation from the mean for six biological replicates; p-values were calculated using a two-tailed student t-test for six biological replicates; “****” = p-value < 0.0001, “***” = 0.0001 < p-value < 0.001, “**” = 0.001 < p-value < 0.01, “*” = 0.01 < p-value < 0.05, “NS” = p-value > 0.05; E) Image of crystal violet-stained wells from plaque assay of infected A549-ACE2 cell supernatant performed with Vero E6 cells; F) Western blots of lysates from A549-ACE2 cells infected with the indicated SARS-CoV-2 variant at an MOI of 0.1 for 24 hours.

## Discussion

In this study, we systematically characterized the p38/MAPK pathway in the context of SARS-CoV-2 infection using a combination of genetics, genomics, and proteomics. We identified the specific members of the p38/MAPK pathway that impact SARS-CoV-2 infection, both negatively and positively, and discovered novel pathway kinase substrates in an unbiased manner. We found that p38β is the major component of the p38/MAPK pathway contributing to SARS-CoV-2 infection and that it does so at a stage of virus replication after viral mRNA synthesis. Additionally, we showed that depletion of p38β leads to a significant induction of IFN-I and pro-inflammatory cytokines. We also identified several phosphosites on the N-terminus of SARS-CoV-2 N that are sensitive to the p38α/β inhibitor, SB203580, and showed that the inhibition of phosphorylation of these residues results in attenuated virus growth and increased IFN-I activity. These findings reveal a novel dynamic between SARS-CoV-2 and p38/MAPK biology, and provide host targets of the p38 pathway that may be leveraged for drug development for the treatment of COVID-19.

### The p38/MAPK pathway activity is significantly increased during SARS-CoV-2 infection in diverse cell types

There are many documented mechanisms by which the p38/MAPK pathway may become activated in the context of viral infection. The PDZ-binding motif of SARS-CoV envelope protein, as well as the SARS-CoV 3a and 7a proteins have been shown to potently activate the p38 pathway, but it is not known whether these activities are conserved for SARS-CoV-2 (30–32). *In vivo*, SARS-CoV-2 infection leads to the production of an inflammatory cytokine signature that includes IL-6 and TNFɑ, which are well documented to activate the p38/MAPK pathway downstream of binding their respective receptors (33). Here, we show that knockdown of several individual MAPKKKs, especially MAP3K8, MAP3K9, and MAP3K11, significantly inhibited SARS-CoV-2 infection. This was surprising as MAPKKKs are generally thought to be functionally redundant.

### p38β is required for SARS-CoV-2 replication in human lung epithelial cells

In the present study, we show that p38β, but not p38ɑ, is required for SARS-CoV-2 replication in human lung epithelial cells. We only analyzed single knockdowns, but it is possible that there are additional components of the p38/MAPK pathway regulating SARS-CoV-2 infection that were masked by functional redundancy in our screen. p38α is presumed to be the major isoform involved in inducing immune responses, as p38β knockout in mice contributes to neither p38-dependent immediate-early gene transcription nor LPS-induced inflammation. However, while immune responses in p38β^-/-^ mice have been assessed in the contexts of LPS stimulation and TNF overexpression, they have not been assessed in the context of viral infection (23). Our findings emphasize the need for further research and reagents to help better characterize p38β. p38β does not appear to be essential as p38β knockout mice are viable and fertile whereas p38ɑ knockout mice are embryonically lethal, and p38β has a distinctly smaller active site than p38ɑ that may allow for the development of specific inhibitors (23). Thus, p38β makes an attractive target for the treatment of COVID-19. Additional studies are needed to determine if p38β is important for the replication of other coronaviruses and other virus families.

### Identification of novel p38β substrates using novel analysis pipeline, KiPPC

We globally quantified changes to the phosphoproteome in response to several perturbations affecting p38/MAPK signal transduction. In combination, these data allowed us to extract log_2_fold-change profiles of annotated p38 substrate phosphosites in response to these perturbations and to identify novel, putative p38β substrates based on the similarity of their profiles to annotated p38 substrate profiles. This <u>ki</u>nase <u>p</u>erturbation <u>p</u>hospho-profile <u>c</u>lustering (KiPPC) method is broadly applicable for kinases for which some annotated substrates have been identified and for which specific inhibitors are available. It could be advantageous to perturb the system with multiple inhibitors, inhibitor concentrations, or timepoints to separate off- and on-target inhibitor effects and to develop more specific substrate profiles to feed into the clustering algorithm. One limitation of this approach is that it cannot determine whether the substrates identified were direct or indirect substrates of the kinase of interest.

As for the substrates we identified, we observed that knockdown of a vast majority of the substrates tested (22/29) resulted in increased SARS-CoV-2 replication, suggesting that these substrates have antiviral activity with respect to SARS-CoV-2 and that their phosphorylation by p38/MAPKs may inactivate their antiviral functions. However, we have not yet tested the ability of phosphorylation site mutants to rescue these effects, which would more conclusively determine their contributions but is beyond the scope of this study. Future work will determine the contributions of individual phosphorylation sites and answer whether p38/MAPKs exert their effects on SARS-CoV-2 infection via a small number substrate with strong effects or the combined impact of many substrates with moderate effects.

In support of these findings, many of these novel substrates have been implicated as relevant in the context of SARS-CoV-2 infection; single nucleotide polymorphisms (SNPs) in their respective phosphosites are naturally occurring in the human population and SNPs have been associated with SARS-CoV-2 disease outcome (TRIM28, ACIN1, TNKS1BP1, HSPB1, and LARP7) (34). Additionally, TRIM28 deficiencies have even been correlated with severe pediatric COVID-19 cases (35).

### SARS-CoV-2 N contains phosphosites sensitive to the p38α/β inhibitor, SB203580

Throughout the virus life cycle, coronavirus N protein performs many crucial functions including oligomerization along the length of viral RNA for protection, enhancing viral polymerase activity, modulating template switching, and innate immunity evasion (36). N post-translational modifications have been documented to affect N activities; avian *Gammacoronaviru*s infectious bronchitis virus N phosphorylation increases the affinity of N for viral RNA compared to non-viral RNA (37). Additionally, *Betacoronavirus* murine hepatitis virus (MHV) N phosphorylation by GSK-3 regulates synthesis of genomic RNA versus subgenomic RNA by promoting template read-through (38). Specific to SARS-CoV-2, many studies have implicated SARS-CoV-2 N phosphorylation in processes including liquid-liquid phase separation and innate immunity activation, but mechanisms to explain phenotypes have yet to be elucidated (39, 40). In this study, we identified phosphosites on SARS-CoV-2 N that are sensitive to a p38 inhibitor. As N phosphorylation is known to affect its activity, it is plausible that p38-dependent N phosphorylation is responsible for the phenotypes we observed, but we cannot exclude the possibility that p38-dependent phosphorylation of host protein(s) may play a more significant role in driving these phenotypes.

## Methods

### Cells

A549 cells, a human lung epithelial cell line (A549, ATCC®, CCL-185), HEK 293T cells, a human kidney epithelial cell line (HEK 293T/17, ATCC®, CRL-11268), and Vero E6 cells (Vero 76, clone E6, Vero E6, ATCC® CRL-1586), an African Green Monkey kidney epithelial cell line, were authenticated by ATCC. A monoclonal ACE2-expressing A549 cell line (A549-ACE2) was a kind gift from Brad Rosenberg. Monoclonal ACE2-expressing STAT1-knockout A549 cells (A549-ACE2ΔSTAT1) were generated as previously described (41, 42). All cell lines were cultured under humidified 5% CO_2_ conditions in 10% v/v fetal bovine serum (FBS, Thermo Fisher Scientific) and 100 I.U. penicillin and 100 µg/mL streptomycin (Pen/Strept, Corning) in Dulbecco’s Modified Eagle Medium (DMEM, Corning). Cells were confirmed negative for mycobacteria monthly (Lonza).

### Viruses

SARS-related coronavirus 2 (SARS-CoV-2) isolate USA-WA1/2020 (NR-52281) was obtained from BEI Resources, NIAID, NIH. Recombinant SARS-CoV-2 (rSARS-CoV-2), based on (isolate USA-WA1/2020), and rSARS-CoV-2^N4A^ (S/T to A mutations at S21, S23, T24, and S26 on SARS-CoV-2 N) were generated as previously described (Ye et al., 2020). Virus stocks were grown on Vero E6 cells by infecting in infection media (2% FBS, Pen/Strept, in DMEM) at an MOI of 0.01. Supernatant was collected 30h post-infection and concentrated through a 100kDa centrifugal filter unit (Amicon). Concentrated virus was washed thrice in phosphate-buffer saline (PBS) and concentrated with a 100kDa centrifugal filter unit (43). Virus stock titers were determined by plaque assay on Vero E6 cells. All work with live virus was done in the CDC/USDA-approved biosafety level 3 (BSL-3) facility of the Icahn School of Medicine at Mount Sinai or the NYU Grossman School of Medicine in accordance to their respective guidelines for BSL-3 work.

### Cell treatment prior to harvest for mass spectrometry

*SB203580 pre-treatment treatment:* three plates each of 2×10^7^ A549-ACE2 cells in 15cm plate format were treated with a final concentration of 1uM SB203580 or 10uM SB203580, and six plates treated with an equal volume of DMSO as the drug treated plates, in 25mL of infection media, and were incubated for one hour. All plates except three DMSO-treated plates (Mock-infected), were infected by adding SARS-CoV-2 at an MOI 0.1 directly to the drug-containing infection media. At 24 hours post-infection, all cells were lysed in urea lysis buffer as described below. *Terminal SB203580 treatment:* Eight plates of 2×10^7^ A549-ACE2 cells in 15cm plate format were infected with SARS-CoV-2 at an MOI 0.1 in 25mL infection media. An additional four plates were mock-infected in 25mL infection media. At 20 hours post-infection, half of the infected replicates were treated with SB203580 (Cell Signaling) in DMSO at a final concentration of 10μM and the other half of the infected replicates and all of the mock-infected replicates were treated with an equal volume of DMSO. 4 hours after drug treatment, all cells were lysed in urea lysis buffer as described below. *siRNA knockdown:* four plates each of 2×10^7^ A549-ACE2 cells in 15cm plate format were transfected with pooled siRNAs against *MAPK11* or *MSK2* (Dharmacon), and eight plates were transfected with pooled siRNAs against NTC (Dharmacon) according to the manufacturing protocol for RNAiMAX (ThermoFisher Scientific). At 48 hours post-transfection, the media on all plates was replaced with 25mL of infection media. All plates except four NTC plates (Mock-infected), were infected by adding SARS-CoV-2 at an MOI 0.1 directly to the drug-containing infection media. At 36 hours post-infection, all cells were lysed in urea lysis buffer as described below.

### Cell lysis and digestion for mass spectrometry

Cells were washed twice in PBS. Cells were lysed in urea lysis buffer containing 8M urea, 100mM ammonium bicarbonate (ABC), 150mM NaCl, protease inhibitor and phosphatase inhibitor cocktail (Thermo Fisher Scientific). Lysates were probe-sonicated on ice 3×1s at 50% power, with 5s of rest in between pulses. Protein content of the lysates was quantified using a micro BCA assay (Thermo Fisher Scientific). 1mg of protein per sample was treated with (Tris-(2-carboxyethyl)phosphine (TCEP) to a final concentration of 4mM and incubated at room temperature (RT) for 30 minutes. Iodoacetamide (IAA) was added to each sample to a final concentration of 10mM, and samples were incubated in the dark at RT for 30 minutes. IAA was quenched by dithiothreitol (DTT) to a concentration of 10mM and incubated in the dark at RT for 30 minutes. Samples were then diluted with 5 sample volumes of 100mM ABC. Trypsin Gold (Promega) was added at a 1:100 (enzyme:protein w/w) ratio and lysates were rotated for 16 hours at RT. 10% v/v trifluoroacetic acid (TFA) was added to each sample to a final concentration of 0.1% TFA. Samples were desalted under vacuum using Sep Pak tC18 cartridges (Waters). Each cartridge was first washed with 1mL 80% acetonitrile (ACN)/0.1% TFA, then with 3×1mL of 0.1% TFA in H_2_O. Samples were then loaded onto cartridges. Cartridges were washed with 3×1mL of 0.1% TFA in H_2_O. Samples were then eluted with 1mL 40% ACN/0.1% TFA. 20μg of each sample was kept for protein abundance measurements, and the remainder was used for phosphopeptide enrichment. Samples were dried by vacuum centrifugation. Protein abundance samples were resuspended in 0.1% formic acid (FA) for mass spectrometry analysis.

### Phosphopeptide enrichment for mass spectrometry

For each sample batch and under vacuum, 500μL (30μL per sample) of 50% Ni-NTA Superflow bead slurry (QIAGEN) was added to a 2mL bio-spin column. Beads were washed with 3×1mL HPLC H_2_O, incubated with 4×1mL 100mM EDTA for 30s, washed with 3× 1mL HPLC H_2_O, incubated 4×1mL with 15mM FeCl_3_ for 1 minute, washed 3×1mL HPLC H_2_O, and washed once with 1mL of 0.5% v/v FA to remove residual iron. Beads were resuspended in 750μL of 80% ACN/0.1% TFA. 1mg of digested peptides were resuspended in 83.33μL 40% ACN/0.1% TFA and 166.67μL 100% ACN/0.1% TFA and 60μL of the bead slurry were added to each sample and incubated for 30 minutes while rotating at RT. A C18 BioSPN column (Nest Group), centrifuged at 110xg for 1 minute each step, was equilibrated with 2×200μL 80% ACN/0.1% TFA. Beads were loaded onto the column and washed with 4×200μL 80% ACN/0.1% TFA, then washed 3×200μL 0.5% FA. Then, 2×200μL 500mM potassium phosphate buffer pH 7 was added to the column and incubated for one minute. Then, 3×200μL 0.5% FA were added to the column. Phosphopeptides were eluted with 2×100μL of 40% ACN/0.1% FA and vacuum centrifuged to dryness. Phosphopeptides were resuspended in 25μL 4% FA/3% ACN for mass spectrometry analysis.

### Mass spectrometry data acquisition

All samples were analyzed on an Orbitrap Eclipse mass spectrometry system (Thermo Fisher Scientific) equipped with an Easy nLC 1200 ultra-high pressure liquid chromatography system (Thermo Fisher Scientific) interfaced via a Nanospray Flex nanoelectrospray source. For all analyses, samples were injected on a C18 reverse phase column (30cm × 75μm (ID)) packed with ReprosilPur 1.9μm particles). Mobile phase A consisted of 0.1% FA, and mobile phase B consisted of 0.1% FA/80% ACN. Peptides were separated by an organic gradient from 5% to 35% mobile phase B over 120 minutes followed by an increase to 100% B over 10 minutes at a flow rate of 300nL/minute. Analytical columns were equilibrated with 6μL of mobile phase A. To build a spectral library, samples from each set of biological replicates were pooled and acquired in data dependent manner. Protein abundance samples were fractionated with Field Assymetric Ion Mobility Spectrometry (FAIMS) fractionation with a FAIMS Pro device (Thermo Fisher Scientific). Each pooled sample was analyzed four times with four FAIMS compensation voltages (CV) (−40V,−55V,−65V,−75V). Data dependent analysis (DDA) was performed by acquiring a full scan over a m/z range of 375-1025 in the Orbitrap at 120,000 resolving power (@200 m/z) with a normalized AGC target of 100%, an RF lens setting of 30%, and an instrument controlled ion injection time. Dynamic exclusion was set to 30 seconds, with a 10ppm exclusion width setting. Peptides with charge states 2-6 were selected for MS/MS interrogation using higher energy collisional dissociation (HCD) with a normalized HCD collision energy of 28%, with three seconds of MS/MS scans per cycle. Similar settings were used for data dependent analysis of phosphopeptide-enriched pooled samples, with a Dynamic Exclusion of 45 seconds and no FAIMS fractionation. Data-independent analysis (DIA) was performed on all individual samples. An MS scan at 60,000 resolving power over a scan range of 390-1010 m/z, an instrument controlled AGC target, an RF lens setting of 30%, and an instrument controlled maximum injection time, followed by DIA scans using 8 m/z isolation windows over 400-1000 m/z at a normalized HCD collision energy of 28%.

### siRNA knockdown

2×10^4^ A549-ACE2 cells in a 96-well plate format were transfected (nine technical replicates), with 1pmol ON TARGETplus siRNA pools (Dharmacon) prepared in 10μL/replicate Opti-MEM (Corning) with a 1:3 ratio of siRNA:RNAiMax (Thermo Fisher Scientific). 48 hours post-transfection, cells were infected with SARS-CoV-2 at a MOI of 0.1 or 2 in infection media. 8-, 30-, or 36-hours post-infection, supernatants were saved for plaque assay, one-third of replicates were fixed in 5% paraformaldehyde (PFA) in PBS for 24 hours, one-third of replicates were lysed in RIPA buffer with SDS (50mM Tris HCl, 150mM sodium chlorate, 1% v/v Triton X-100, 0.5% v/v sodium deoxycholate, 1% w/v sodium dodecyl sulfate, and protease inhibitors (MilliporeSigma)) and saved for Western blotting, and the last third of replicates were lysed in RLT buffer (QIAGEN) and saved for RNA extraction and RT-qPCR analysis.

### Immunofluorescence assay

Fixed cells were washed with PBS and permeabilized for 10 minutes in 0.2% v/v Triton X-100 in PBS. Cells were incubated in blocking buffer (3% w/v BSA, 0.1% v/v Triton X-100, 0.2% w/v fish gelatin in PBS) at room temperature for one hour. Cells were incubated in primary antibody (1:1000 mouse anti-SARS-CoV-1/2 N 1C7C7 antibody, a kind gift from Thomas Moran) in antibody buffer (1% w/v BSA, 0.03% v/v Triton X-100, 0.1% fish gelatin in PBS) overnight at 4°C. Cells were washed thrice in PBS. Cells were incubated in 1:1000 anti-mouse AlexaFluor488 or anti-mouse AlexaFluor594 (Thermo Fisher Scientific) and 4’,6-diamidino-2-phenylindole counterstain (DAPI, Thermo Fisher Scientific) in antibody buffer at room temperature for one hour. Cells were washed thrice in PBS and imaged in 100μL PBS on a Celígo Imaging Cytometer (Nexcelcom Bioscience) or a CX7 Imaging Cytometer (ThermoFisher Scientific). Celígo software or CX7 software was used to quantify the total number of cells by DAPI nuclear staining and the number of N-positive cells by N staining.

### Plaque assay

25μL of virus-containing supernatant was serially 10-fold diluted in infection media. 100μL inoculum was added to confluent Vero E6 cells in a 24-well plate format and incubated at room temperature for one hour and agitated often to avoid drying. 1mL semi-solid overlay (0.1% w/v agarose, 4% v/v FBS, and Pen/Strept in DMEM) was added to each well and plates were incubated at 37C for 48 hours. Cells were fixed in 5% PFA in PBS for 24 hours at room temperature. Cells were washed twice with water. Cells were incubated in 0.5mL staining dye (2% w/v crystal violet, 20% v/v ethanol in water) for 10 minutes at room temperature. Stained cells were washed with water and allowed to dry before plaques were counted and plaque-forming units per mL was calculated as follows: ((# of plaques)/(mL of inoculum*10^dilution^ ^factor^)).

### Western blotting

Lysates were incubated at 95C for 10 minutes in Laemmli sample buffer (Bio-Rad Laboratories). Lysates were run on an SDS-PAGE gel with a protein ladder standard (Bio-Rad Laboratories) and transferred to a nitrocellulose membrane (Bio-Rad Laboratories). Blots were incubated in 5% milk in TBST for one hour at room temperature. Blots were incubated in primary antibody in 1% milk in TBST overnight at 4°C. Blots were washed thrice for 5 minutes in TBST. Blots were incubated in secondary HRP-conjugated (Bio-Rad Laboratories) or infrared-conjugated secondary antibodies (LICOR Biosciences) in 1% milk in TBST. Blots were washed thrice for 5 minutes in TBST. Blots were imaged on a Chemiluminescence digital imager (Bio-Rad Laboratories) using FEMTO ECL reagent (Thermo Fisher Scientific) or an infrared digital imager (LICOR Biosciences).

### Cell viability assay

2×10^4^ A549-ACE2 cells in a 96-well white-bottom plate were transfected in triplicate with siRNA pools. 72 hours post-transfection, the plate was equilibrated to room temperature for 30 minutes. 100μL of Titerglo buffer with substrate was added to each well (Promega Corporation). Plate was nutated at room temperature for 2 minutes to lyse the cells. Plate was incubated at room temperature for 10 minutes. Plate was read for luminescence end-point kinetics with a 1s integration on a Cytation 5 Plate Reader using Gen5 software (Biotek Instruments). Relative luminescence units for each siRNA condition were normalized to NTC siRNA and displayed as a percentage.

### Entropy of N analysis

Entropy values for each amino acid on N for SARS-CoV-2 sequences from GISAID were downloaded from Next Strain on December 13, 2021 (Hadfield et al., 2018). mRNA sequencing and analysis 2×10^5^ A549-ACE2 cells were transfected with siRNA pools as previously indicated. 48 hours post-transfection, cells were infected with SARS-CoV-2 at an MOI of 0.75 in infection media. 8 hours post-infection, cells were lysed in 1mL of Trizol (Invitrogen). RNA was extracted and DNase I treated using Direct-zol RNA Miniprep kit (Zymo Research) according to the manufacturer’s protocol. RNA-seq libraries of polyadenylated RNA were prepared with the TruSeq RNA Library Prep Kit v2 (Illumina) according to the manufacturer’s protocols. Libraries were sequenced with an Illumina NextSeq 500 platform. Raw sequencing reads were aligned to the hg19 human genome with the Basespace RNA-Seq Aligment application (Illumina). GO-term enrichment was performed using Biojupies (44). Alignment to viral genomes was performed using bowtie2 (45). The SARS-CoV-2 USA/WA1/2020 strain genome was used for analysis in this study (GenBank: MN985325). Gene set enrichment analysis was performed with the enrichr package in R (46). All raw and processed mRNA-Seq data is available on NCBI GEO GSE183999.

### RNA extraction and RT-qPCR analysis

Cells were lysed in RLT buffer and RNA was extracted using a RNeasy 96 kit (QIAGEN) according to the manufacturer’s protocol. 1-step RT-qPCR was performed on 2μL of RNA using the Luna® Universal One-Step RT-qPCR Kit (NEB Biosciences) and primers for β-tubulin (F: 5’-GCCTGGACCACAAGTTTGAC-3’; R: 5’-TGAAATTCTGGGAGCATGAC-3’), SARS-CoV-2 *NSP14* (F: 5’-TGGGGYTTTACRGGTAACCT-3’; R: 5’-AACRCGCTTAACAAAGCACTC-3’), and *TRS-N* (F: 5’-CTCTTGTAGATCTTCTCTAAACGAAC-3’; R: 5’-GGTCCACCAAACGTAATGCG-3’) as previously described (41, 47). Reactions were analyzed on a Lightcycler 480 II Instrument (Roche). ΔΔ cycle threshold values were calculated relative to mock-infected samples and NTC samples.

### Statistical analysis

All experiments were performed in at least biological triplicate with at least three technical replicates per biological replicate, when appropriate. Biological replicates are defined here as separate wells or plates of an experiment. Technical replicates are defined here as separate instrumental measurements within single biological replicate. All experiments, with the exception of the mass spectrometry and RNA-Seq experiments due to the extensive sample processing, were performed at least separate times with separate passages of cells on separate days to ensure results were consistent. Unless otherwise noted, error bars indicate one standard deviation from the mean of three biological replicates. Unless otherwise noted, one-way ANOVA tests with post hoc testing by Tukey’s method (in comparison to a control) were performed to generate p-values in R with the rstatix package (Kassambara 2019). “****” = p-value < 0.0001, “***” = 0.0001 < p-value < 0.001, “**” = 0.001 < p-value < 0.01, “*” = 0.01 < p-value < 0.05, “NS” = p-value > 0.05.

### Mass spectrometry data analysis

All raw mass spectrometry data generated in this study were analyzed by the Spectronaut software suite (Biognosys) (48). DDA-MS data were analyzed to build a spectral library by searching against a protein sequence database comprised of SwissProt human sequences (downloaded on October 10, 2019) and SARS-CoV-2 strain USA/WA1/2020 sequences using Biognosys factory settings, which considered a static modification for cysteine carbamidomethylation and variable modifications for methionine oxidation and protein N-terminal acetylation. We added variable modifications for serine/threonine/tyrosine phosphorylation for phospho-enriched samples. All DDA-MS runs generated in this study were combined to make one spectral library against which all DIA-MS data were analyzed. DIA-MS data were also analyzed by Spectronaut to extract fragment ion intensity information based on the spectral library described above using Biognosys factory settings. The data were exported as a tab-delimited, MSstats-formatted report. Spectronaut reports were analyzed by the MSstats package in the Rstudio statistical programming environment (49). Before MSstats analysis, protein group accessions were converted to phosphosite group accessions with a Perl script. Data were processed by MSstats to equalize medians, summarize features using Tukey’s median polish, and impute missing values by accelerated failure model. Intensities below the 0.999 quantile were considered missing values. Principal component analysis was performed on MSstats estimated intensities using the prcomp function in R and the first two principal components plotted for all data sets. Sample correlation analysis was performed by pairwise Pearson correlation coefficient calculation in R.

### Gene ontology (GO) enrichment and kinase activity analyses

GO enrichment and kinase activity analyses were performed using the GSEA method with the fgsea package in R (Subramanian et al., 2005). For kinase activity analysis, kinase substrate interactions were derived from the PhosphositePlus Kinase Substrate Dataset (21). Protein substrates of each kinase were compiled into gene sets. For gene ontology enrichment analysis, GO terms and definitions were downloaded from the gene ontology resource (downloaded on February 18, 2021) and genes within each GO term were grouped together as gene sets. For both Go enrichment analysis and kinase activity analysis, for each comparison considered, the data were ranked by log_2_(fold-change) subjected to fgsea testing using the gene sets described above.

### Unbiased identification of p38β substrates

For hierarchical clustering, comparisons indicated in Figure 5 were first filtered for missing log-fold-change values in any comparison. Distances were calculated based on Euclidean distance using the dist function in R and the data were hierarchically ordered using the hclust function in R. Next, the data were divided into *n* clusters in decreased order of dendrogram height in 99 iterations with *n* ranging from 2 to 100 using the cutree function in R. For each iteration, the enrichment of annotated p38α/β substrates within each cluster was calculated using a hypergeometric test with the dhyper function in R. p-values were adjusted by Benjamini-Hochberg method with the p.adjust function in R. Annotations of p38α/β substrates were derived from the Phosphosite Plus Kinase Substrate data set (21). phosphosites in the cluster with the minimum hypergeometric p-value in all iterations across all clusters comprises our putative p38β substrates.

## Data availability

Raw mass spectrometry data have been deposited to the ProteomeXchange Consortium via the PRIDE partner repository with the dataset identifier PXD035451. For the purposes of manuscript review, the data can be accessed with the username and password LMEfkIpz.

## Acknowledgments

We would like to acknowledge the following funding sources: NIH R01 AI151029, NIH 0255-E382 (BRR), NIH U01 AI150748 (BRR), NIH R21AI164043 (JRJ), and NIH U19AI118610 (JRJ). We would also like to thank Dr. Ludovic P. Desvignes, Dr. Dominic Papandrea, and Dr. Randy Albrecht, for all their support in regard to working in the BSL-3 facilities.

## Author Contributions

Conceptualization: CAH, JRJ

Proteomics sample preparation: CAH, APK, PA Proteomics data acquisition: APK Recombinant virus generation: CY, CAH

Cell line generation: OD Experiments: CAH

RNA-seq preparation and acquisition: MP RT-qPCR: IG

Kinase substrate motif analysis: TY Data analysis: JRJ, CAH

Figure generation: JRJ, CAH Manuscript Preparation: JRJ, CAH

Manuscript editing: BRR, OD, BRT, BENP, CAH, JRJ Literature Review: JRJ, CAH

Work Supervision: BRR, LCC, LMS, BRT, JRJ

## Supplemental Table Legends

Table S1: Table of processed LC-MS data. Values in wide format (easier to view, labeled on tab) with column headings: accession number (with phosphorylated residue), condition, log_2_(fold-changes), p-values, adjusted p-values, protein name, and protein description, and each row is a protein group or phosphosite group detected; or long format (easier to analyze, labeled on tab) with column headings: accession number (with phosphorylated residue), protein name, protein description, and log_2_(fold-changes), p-values, and adjusted p-values for each indicated comparison. Tabs are separate experiments and annotated either protein abundance (AB) or phospho-enriched (PH), and either wide format, or long format. Data are available via ProteomeXchange with identifier PXD035451.

Table S2: Table of phospho-enriched MS data from this paper and several publications comparing SARS-CoV-2-infected cells to mock-infected cells after 24h (Bojkova et al., 2020; Bouhaddou et al., 2020; Hekman et al., 2020; Stukalov et al., 2021). Rows indicate each detected phosphorylation group and column headings are accession number (with phosphorylated residue), each dataset condition, “Nup” *N* number of datasets the site is upregulated in, and “Ndown” is *N* number of datasets the site is downregulated in, protein name, and protein description.

Table S3: Gene set enrichment analysis results for kinase activity including kinase of interest (“pathway”), p-values, adjusted p-values, enrichment scores (“ES”), normalized enrichment scores (“NES”), number of substrates contributing (“size”), substrates contributing (“leadingEdgeText”), for each dataset comparison of infected compared to mock (see also Figure 1F and S1B).

Table S4: Gene set enrichment analysis results for gene ontology (GO) comparing each dataset infected over mock-infected log_2_(fold-changes), with column headings: GO term (“pathway”), p-values, adjusted p-values, enrichment scores (“ES”), normalized enrichment scores (“NES”), number of proteins contributing (“size”), corresponding dataset, GO term definition, proteins contributing (“leadingEdgeText”), and type of MS analysis (protein abundance or phosphorylation) (see also Figure S1C).

Table S5: Table of processed mRNA-sequencing data with the column headings: gene name (“X”), log_2_(fold-change) (“logFC”), p-value, adjusted p-value, and condition comparison.

Table S6: Table of GO terms from enrichr analysis with the column headings: GO term description with term (“Term”), number of genes associated with enrichment (“x”), number of genes total in the term (“size”), p-value, adjusted p-value, genes associated with enrichment (“Genes”), condition comparison, and “direction” if terms is enriched by upregulated (UP) or downregulated (DOWN) genes (see also Figure 3E).

Table S7: Table of GO enrichment analysis of differential gene expression results from MS experiment comparing siRNA-perturbed conditions with column headings: GO terms (“pathway”), p-values, adjusted p-values, enrichment scores (“ES”), normalized enrichment scores (“NES”), total number of genes associated with the GO term “size”, condition comparison, and GO term definition (see also Figure 4D).

Table S8: Table of p38 substrate “cluster of interest” phosphosite groups with column headings: accession number with phosphorylated residue, protein name, condition comparison log_2_(fold-changes), and annotation if known p38 substrate, + = yes, - = no (see also Figure 5E-H).

---

**Figure S1:**
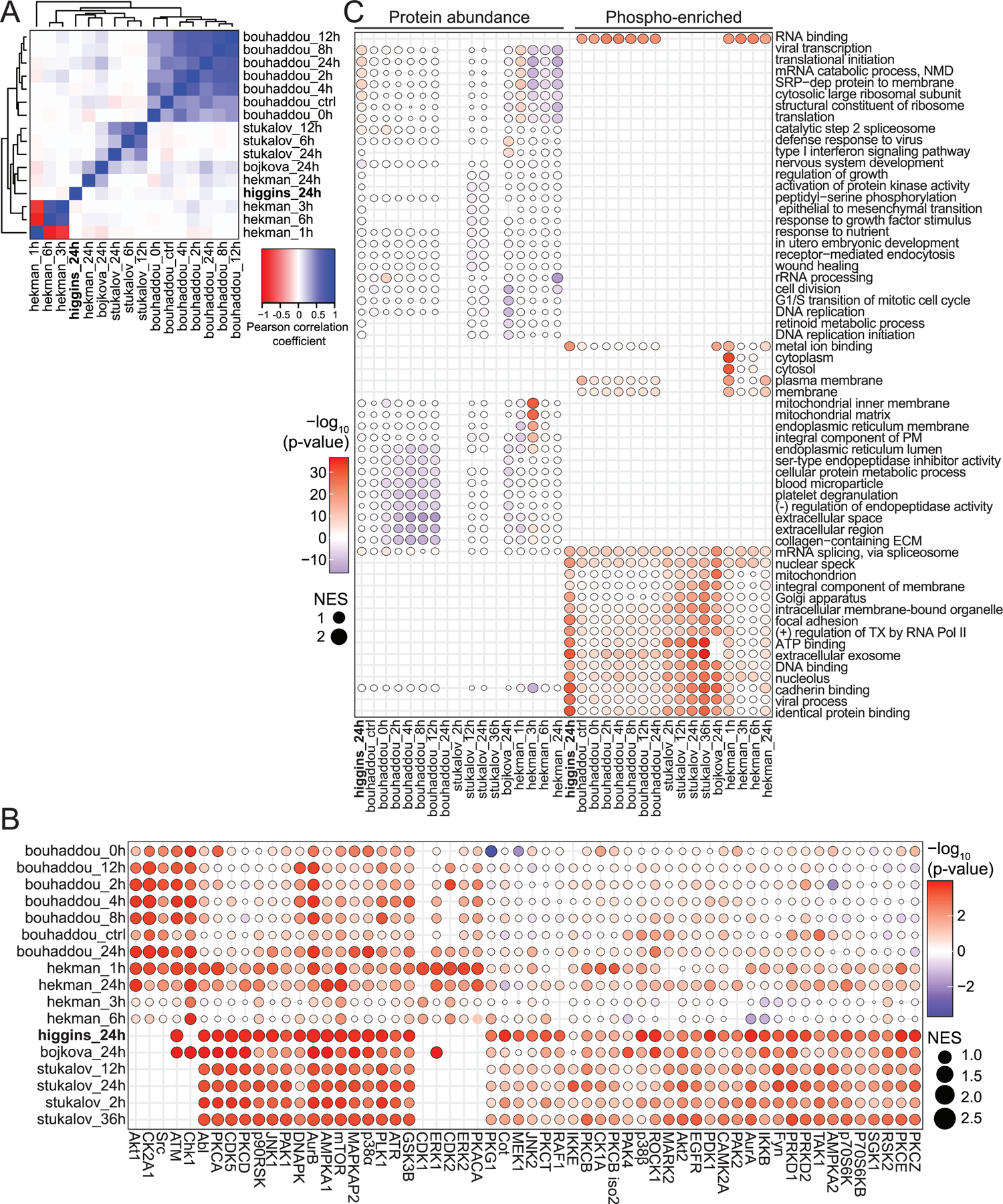
A) Heatmap of pairwise Pearson coefficients for protein group log_2_(fold-change) profiles from this study and published studies indicated; B) Bubble plot of kinase activity analysis based on phosphosite group log_2_(fold-change) profiles from this study and published studies indicated; the absolute value of the normalized enrichment score (NES) is indicated by node sizes and the -log_10_(p-value) is indicated by the color scale. C) Bubble plot of gene ontology enrichment analysis of protein abundance and phosphosite group log_2_(fold-change) profiles from this study and published studies indicated; the absolute value of the NES is indicated by node sizes and the -log_10_(p-value) is indicated by the color scale.

**Supplemental Figure 2:**
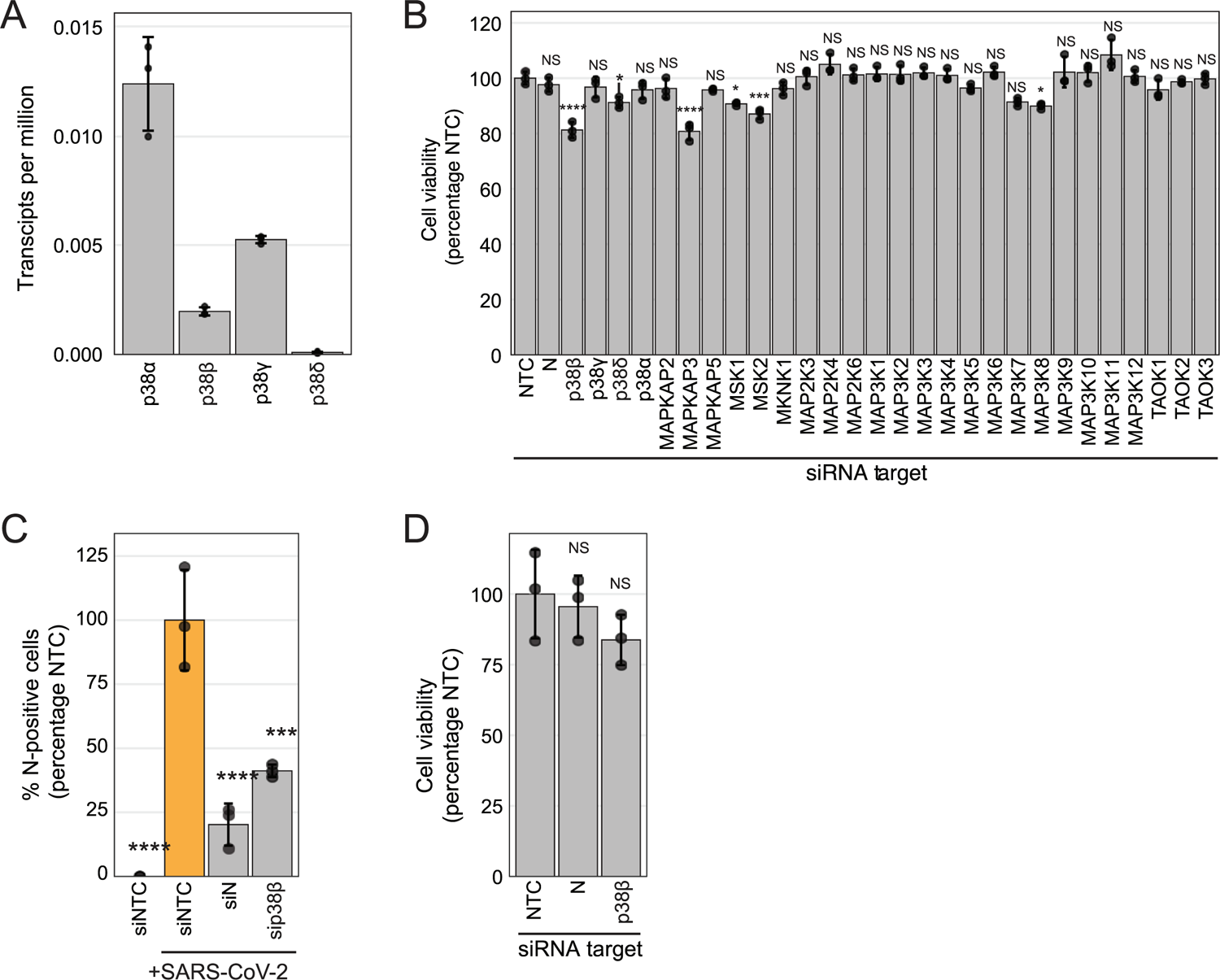
A) Plot of transcripts per million for each indicated p38 isoform from mRNA-sequencing performed on A549-ACE2 cells; B) Plot of A549-ACE2 cell viability normalized to siNTC; C) Plot of the percent of SARS-CoV-2 N-positive cells analyzed using immunofluorescence cytometry, represented as a percentage compared to the infected control condition, for separate Dharmacon ON-TARGETplus siRNA pools targeting NTC or sip38β, compared to Dharmacon siGENOME siRNA pools used in Figure 2B, after SARS-CoV-2 infection at an MOI of 0.1 for 30h in A549-ACE2 cells; D) Plot of A549-ACE2 cell viability normalized to siNTC for siRNA pools used in S2C; error bars represent one standard deviation from the mean for three biological replicates; p-value annotations were calculated using a one-way ANOVA test with post hoc testing using Tukey’s method comparing each condition to siNTC for three biological replicates; “****” = p-value < 0.0001, “***” = 0.0001 < p-value < 0.001, “**” = 0.001 < p-value < 0.01, “*” = 0.01 < p-value < 0.05, “NS” = p-value > 0.05;

**Supplemental Figure 3:**
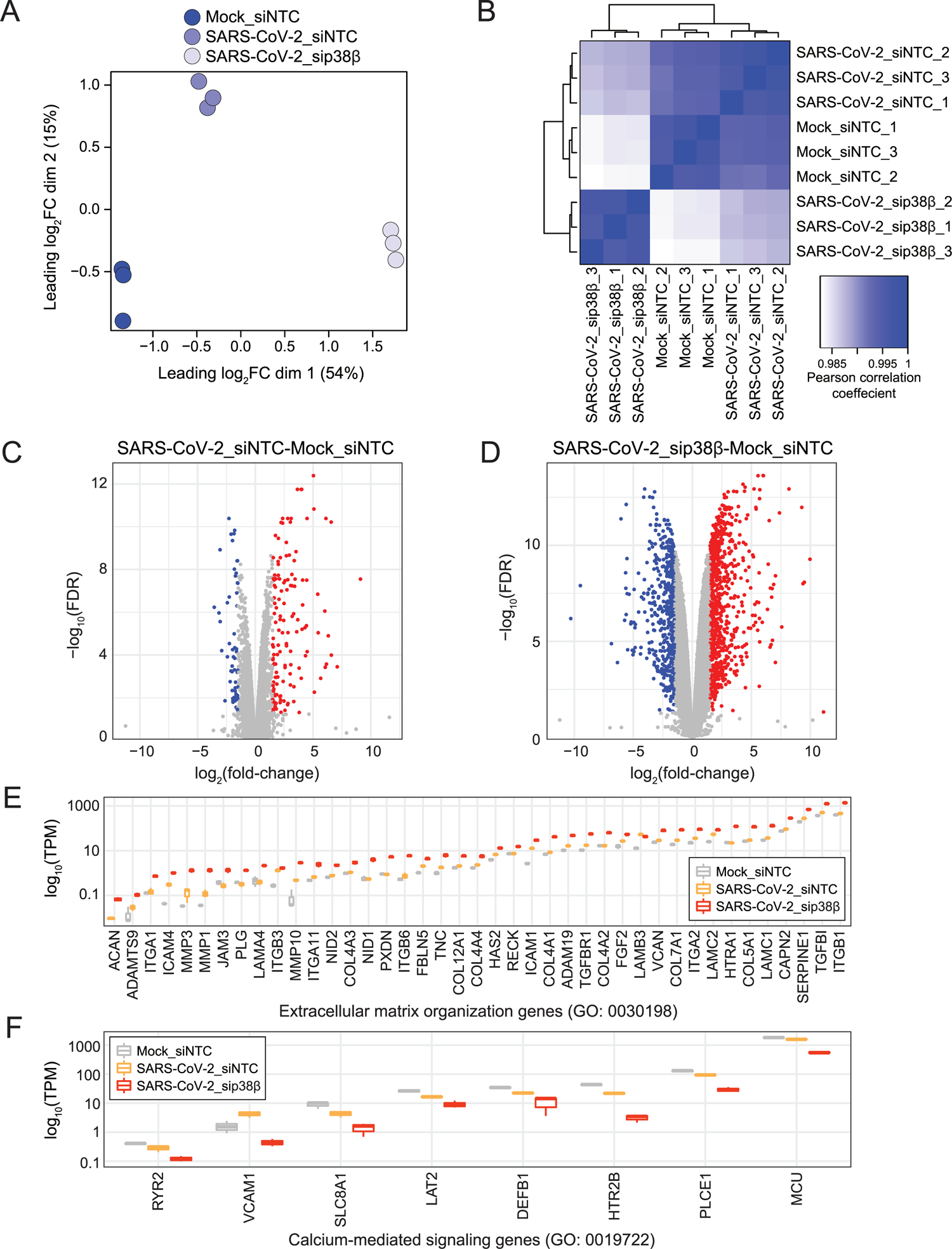
A) Plot of principal component analysis of mRNA-Seq samples; B) Heatmap of Pearson correlation analysis of mRNA-Seq samples; C-D) Volcano plot of differentially expressed genes for the indicated condition comparisons; grey is grey is not differentially expressed, red is upregulated and blue is downregulated; E-F) Plot of log_10_(transcripts per million) for each gene represented in the indicated GO term for each condition, from same analysis as 3D.

**Supplemental Figure 4:**
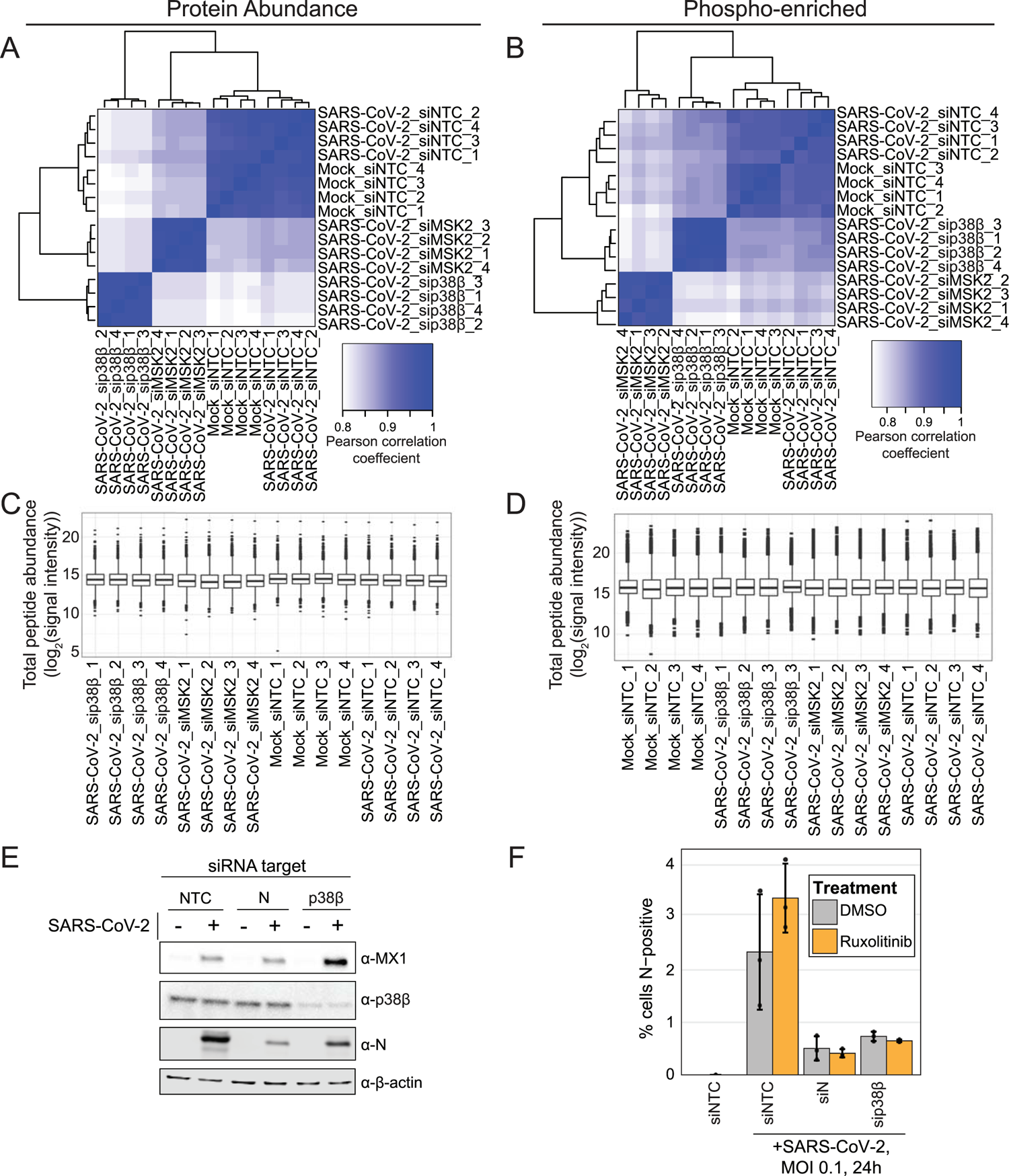
A) Heatmap of Pearson’s correlation analysis of protein abundance mass spectrometry samples; B) Heatmap of Pearson’s correlation analysis of phosphopeptide-enriched mass spectrometry samples; C) Plot of peptide abundance (mass spectrometry signal intensity) for each biological replicate of protein abundance samples; D) Plot of phosphopeptide abundance (mass spectrometry signal intensity) for each biological replicate of phosphopeptide-enriched samples; E) Western blot of lysates from cells transfected with siRNA targeting each indicated gene and infected with SARS-CoV-2 MOI 0.1 or mock-infected for 30h in A549-ACE2 cells; F) Plot of the percent of SARS-CoV-2 N-positive cells analyzed using immunofluorescence cytometry for each indicated transfection condition after SARS-CoV-2 infection at an MOI of 0.1 for 30h in A549-ACE2 cells in the presence of DMSO or ruxolitinib; error bars represent one standard deviation from the mean for three biological replicates; p-values were calculated using a one-way ANOVA test with post hoc testing using Tukey’s method comparing each condition between each cell type for three biological replicates; “****” = p-value < 0.0001, “***” = 0.0001 < p-value < 0.001, “**” = 0.001 < p-value < 0.01, “*” = 0.01 < p-value < 0.05, “NS” = p-value > 0.05.

**Supplemental Figure 5:**
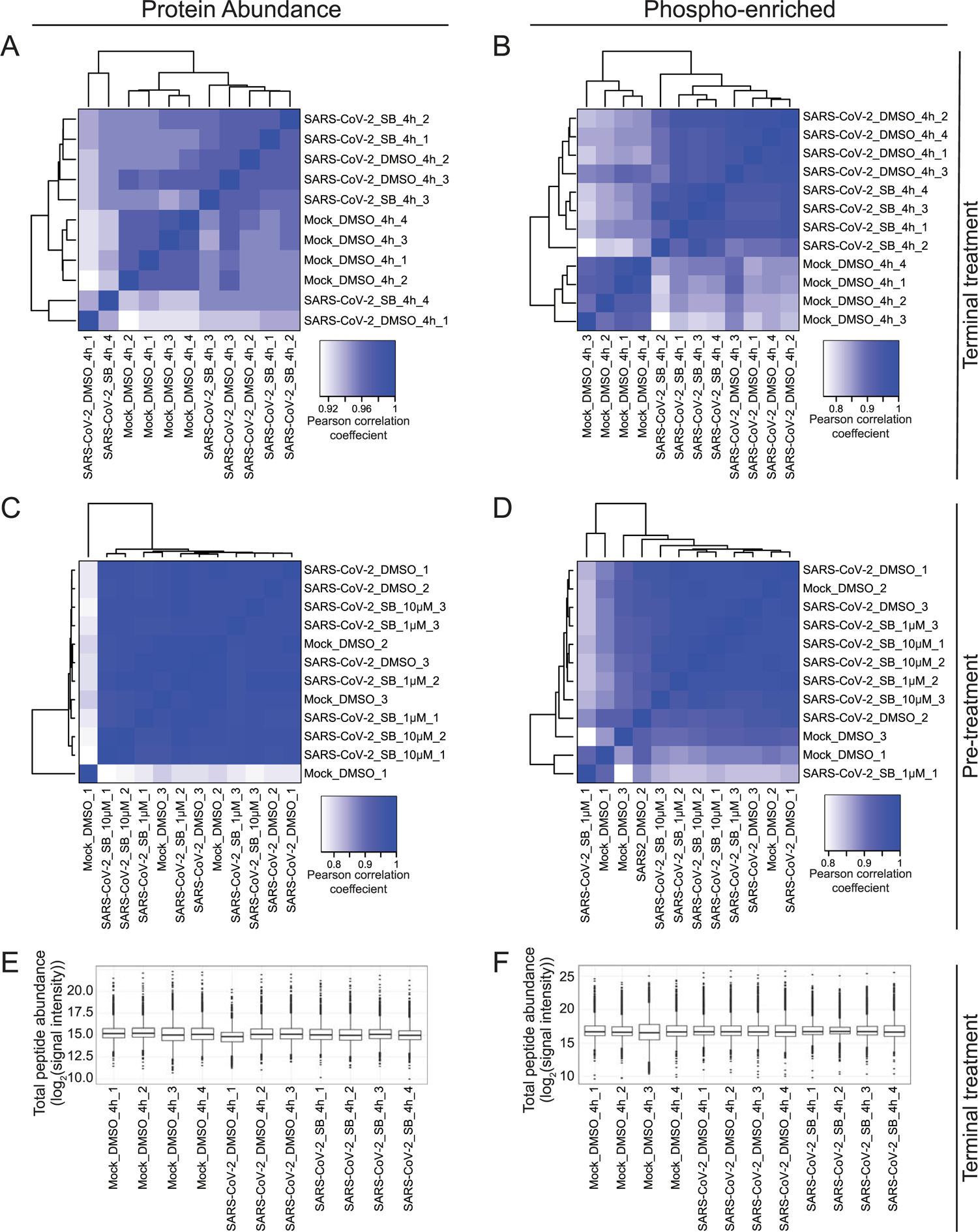
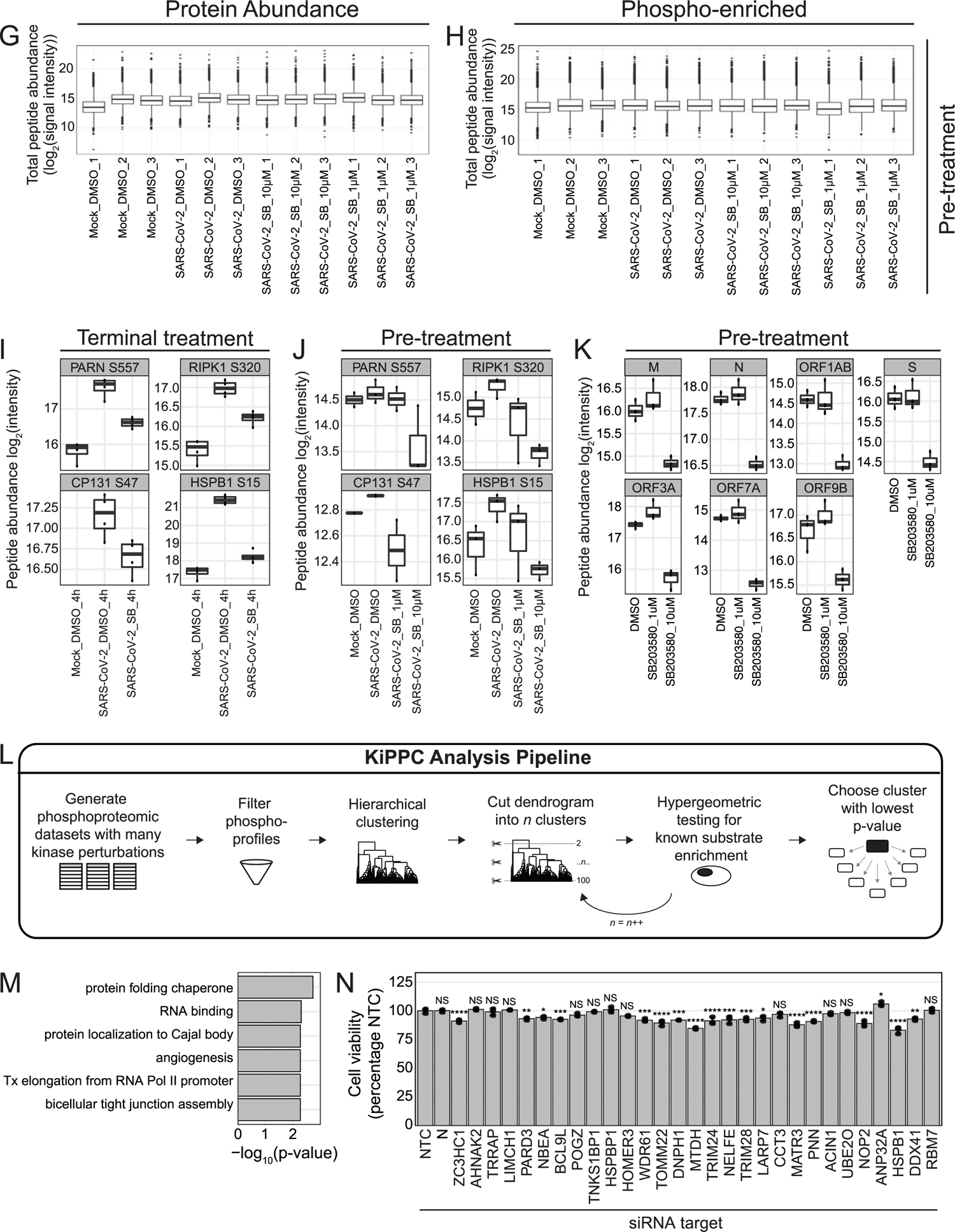
A-B) Heatmap of Pearson’s correlation analysis of protein abundance (A) or phospho-enriched (B) mass spectrometry samples from “terminal treatment” experiment arm; C-D) Heatmap of Pearson’s correlation analysis of protein abundance (A) or phospho-enriched (B) mass spectrometry samples from “pre-treatment” experiment arm; E-F) Plot of peptide abundance (log_2_(mass spectrometry signal intensity)) for each replicate of protein abundance (E) or phospho-enriched (F) samples from “terminal treatment” experiment arm; E-F) Plot of peptide abundance (log_2_(signal intensity)) for each replicate of protein abundance (G) or phospho-enriched (H) samples from “pre-treatment” experiment arm; I) Plot of log_2_(signal intensity)of known p38β substrates from terminal-treatment experiment for each comparison; J) Plot log_2_(signal intensity) of known p38β substrates from pre-treatment experiment for each comparison; K) Plot log_2_(signal intensity) of detected SARS-CoV-2 proteins from pre-treatment experiment for each comparison; L) Schematic of kinase perturbation phospho-profile clustering (KiPPC) pipeline; M) Gene ontology terms enriched from cluster-of-interest proteins using GSEA; N) Plot of A549-ACE2 cell viability normalized to siNTC for each cluster-of-interest-gene siRNA transfection; error bars represent one standard deviation from the mean for three biological replicates; p-values were calculated using a one-way ANOVA test with post hoc testing using Tukey’s method comparing each condition to the infected control condition, for three biologicals replicates; “****” = p-value < 0.0001, “***” = 0.0001 < p-value < 0.001, “**” = 0.001 < p-value < 0.01, “*” = 0.01 < p-value < 0.05, “NS” = p-value > 0.05.

**Supplemental Figure 6:**
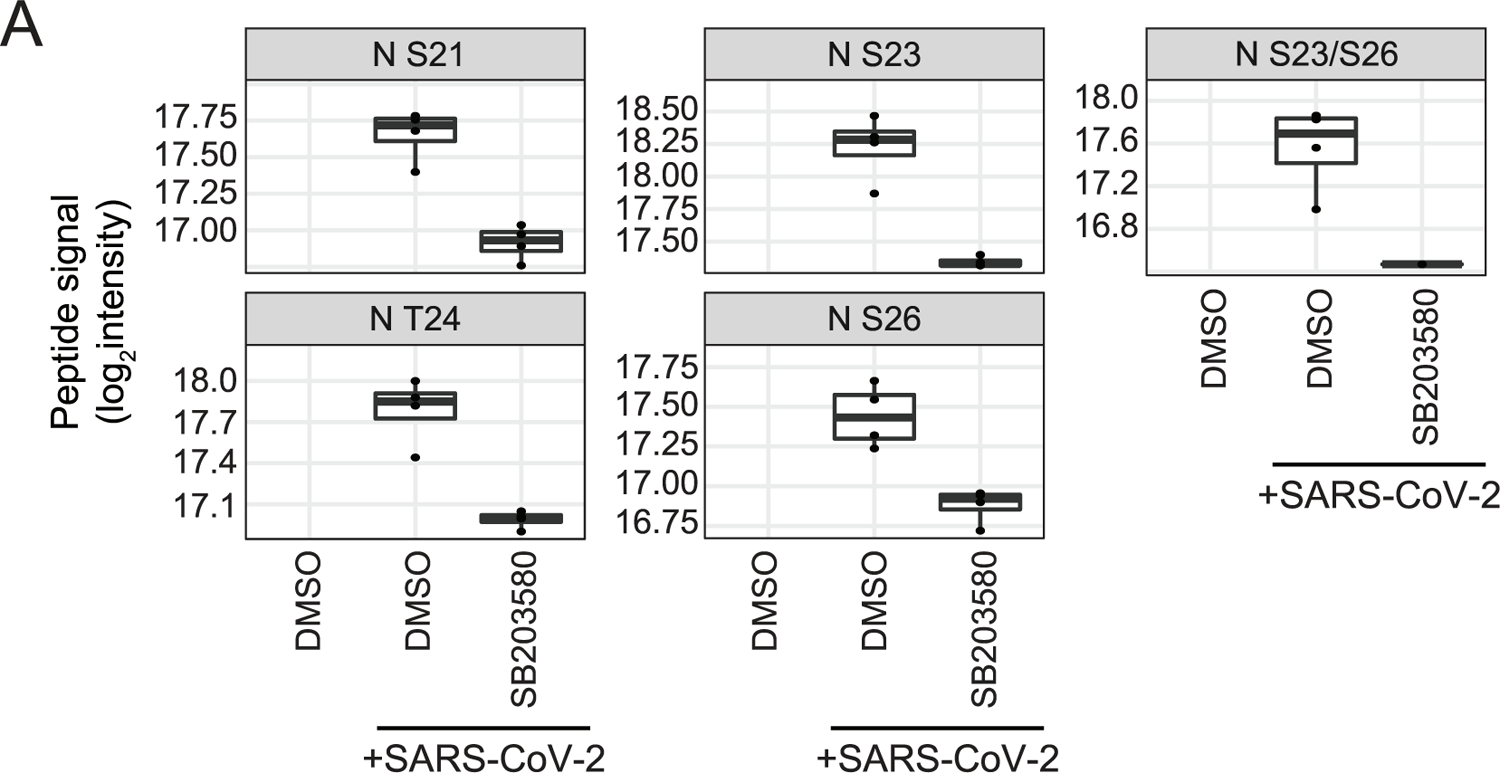
A) Plot of log_2_(signal intensity) of each significantly differentially abundant phosphosite group on SARS-CoV-2 N from the terminal-treatment experiment arm (Figure 5A).

## References

1. Mehta P, McAuley DF, Brown M, Sanchez E, Tattersall RS, Manson JJ, Hlh Across Speciality Collaboration UK. 2020. COVID-19: consider cytokine storm syndromes and immunosuppression. Lancet 395:1033-1034.

2. Merad M, Martin JC. 2020. Pathological inflammation in patients with COVID-19: a key role for monocytes and macrophages. Nat Rev Immunol 20:355–362.

3. Blanco-Melo D, Nilsson-Payant BE, Liu WC, Uhl S, Hoagland D, Moller R, Jordan TX, Oishi K, Panis M, Sachs D, Wang TT, Schwartz RE, Lim JK, Albrecht RA, tenOever BR. 2020. Imbalanced Host Response to SARS-CoV-2 Drives Development of COVID-19. Cell 181:1036–1045 e9.

4. de Vries M, Mohamed AS, Prescott RA, Valero-Jimenez AM, Desvignes L, O’Connor R, Steppan C, Devlin JC, Ivanova E, Herrera A, Schinlever A, Loose P, Ruggles K, Koralov SB, Anderson AS, Binder J, Dittmann M. 2021. A comparative analysis of SARS-CoV-2 antivirals characterizes 3CL(pro) inhibitor PF-00835231 as a potential new treatment for COVID-19. J Virol doi:10.1128/JVI.01819-20.

5. Good SS, Westover J, Jung KH, Zhou XJ, Moussa A, La Colla P, Collu G, Canard B, Sommadossi JP. 2021. AT-527, a Double Prodrug of a Guanosine Nucleotide Analog, Is a Potent Inhibitor of SARS-CoV-2 In Vitro and a Promising Oral Antiviral for Treatment of COVID-19. Antimicrob Agents Chemother 65.

6. Grein J, Ohmagari N, Shin D, Diaz G, Asperges E, Castagna A, Feldt T, Green G, Green ML, Lescure FX, Nicastri E, Oda R, Yo K, Quiros-Roldan E, Studemeister A, Redinski J, Ahmed S, Bernett J, Chelliah D, Chen D, Chihara S, Cohen SH, Cunningham J, D’Arminio Monforte A, Ismail S, Kato H, Lapadula G, L’Her E, Maeno T, Majumder S, Massari M, Mora-Rillo M, Mutoh Y, Nguyen D, Verweij E, Zoufaly A, Osinusi AO, DeZure A, Zhao Y, Zhong L, Chokkalingam A, Elboudwarej E, Telep L, Timbs L, Henne I, Sellers S, Cao H, Tan SK, Winterbourne L, Desai P, et al. 2020. Compassionate Use of Remdesivir for Patients with Severe Covid-19. N Engl J Med 382:2327-2336.

7. Owen DR, Allerton CMN, Anderson AS, Aschenbrenner L, Avery M, Berritt S, Boras B, Cardin RD, Carlo A, Coffman KJ, Dantonio A, Di L, Eng H, Ferre R, Gajiwala KS, Gibson SA, Greasley SE, Hurst BL, Kadar EP, Kalgutkar AS, Lee JC, Lee J, Liu W, Mason SW, Noell S, Novak JJ, Obach RS, Ogilvie K, Patel NC, Pettersson M, Rai DK, Reese MR, Sammons MF, Sathish JG, Singh RSP, Steppan CM, Stewart AE, Tuttle JB, Updyke L, Verhoest PR, Wei L, Yang Q, Zhu Y. 2021. An oral SARS-CoV-2 M(pro) inhibitor clinical candidate for the treatment of COVID-19. Science 374:1586–1593.

8. Sheahan TP, Sims AC, Zhou S, Graham RL, Pruijssers AJ, Agostini ML, Leist SR, Schafer A, Dinnon KH 3rd, Stevens LJ, Chappell JD, Lu X, Hughes TM, George AS, Hill CS, Montgomery SA, Brown AJ, Bluemling GR, Natchus MG, Saindane M, Kolykhalov AA, Painter G, Harcourt J, Tamin A, Thornburg NJ, Swanstrom R, Denison MR, Baric RS. 2020. An orally bioavailable broad-spectrum antiviral inhibits SARS-CoV-2 in human airway epithelial cell cultures and multiple coronaviruses in mice. Sci Transl Med 12.

9. Wahl A, Gralinski LE, Johnson CE, Yao W, Kovarova M, Dinnon KH, 3rd, Liu H, Madden VJ, Krzystek HM, De C, White KK, Gully K, Schafer A, Zaman T, Leist SR, Grant PO, Bluemling GR, Kolykhalov AA, Natchus MG, Askin FB, Painter G, Browne EP, Jones CD, Pickles RJ, Baric RS, Garcia JV. 2021. SARS-CoV-2 infection is effectively treated and prevented by EIDD-2801. Nature 591:451-457.

10. Group RC, Horby P, Lim WS, Emberson JR, Mafham M, Bell JL, Linsell L, Staplin N, Brightling C, Ustianowski A, Elmahi E, Prudon B, Green C, Felton T, Chadwick D, Rege K, Fegan C, Chappell LC, Faust SN, Jaki T, Jeffery K, Montgomery A, Rowan K, Juszczak E, Baillie JK, Haynes R, Landray MJ. 2021. Dexamethasone in Hospitalized Patients with Covid-19. N Engl J Med 384:693–704.

11. Bouhaddou M, Memon D, Meyer B, White KM, Rezelj VV, Correa Marrero M, Polacco BJ, Melnyk JE, Ulferts S, Kaake RM, Batra J, Richards AL, Stevenson E, Gordon DE, Rojc A, Obernier K, Fabius JM, Soucheray M, Miorin L, Moreno E, Koh C, Tran QD, Hardy A, Robinot R, Vallet T, Nilsson-Payant BE, Hernandez-Armenta C, Dunham A, Weigang S, Knerr J, Modak M, Quintero D, Zhou Y, Dugourd A, Valdeolivas A, Patil T, Li Q, Huttenhain R, Cakir M, Muralidharan M, Kim M, Jang G, Tutuncuoglu B, Hiatt J, Guo JZ, Xu J, Bouhaddou S, Mathy CJP, Gaulton A, Manners EJ, et al. 2020. The Global Phosphorylation Landscape of SARS-CoV-2 Infection. Cell 182:685–712 e19.

12. Roux PP, Blenis J. 2004. ERK and p38 MAPK-activated protein kinases: a family of protein kinases with diverse biological functions. Microbiol Mol Biol Rev 68:320–44.

13. Lapointe CP, Grosely R, Johnson AG, Wang J, Fernandez IS, Puglisi JD. 2021. Dynamic competition between SARS-CoV-2 NSP1 and mRNA on the human ribosome inhibits translation initiation. Proc Natl Acad Sci U S A 118.

14. Bojkova D, Klann K, Koch B, Widera M, Krause D, Ciesek S, Cinatl J, Munch C. 2020. Proteomics of SARS-CoV-2-infected host cells reveals therapy targets. Nature 583:469–472.

15. Hekman RM, Hume AJ, Goel RK, Abo KM, Huang J, Blum BC, Werder RB, Suder EL, Paul I, Phanse S, Youssef A, Alysandratos KD, Padhorny D, Ojha S, Mora-Martin A, Kretov D, Ash PEA, Verma M, Zhao J, Patten JJ, Villacorta-Martin C, Bolzan D, Perea-Resa C, Bullitt E, Hinds A, Tilston-Lunel A, Varelas X, Farhangmehr S, Braunschweig U, Kwan JH, McComb M, Basu A, Saeed M, Perissi V, Burks EJ, Layne MD, Connor JH, Davey R, Cheng JX, Wolozin BL, Blencowe BJ, Wuchty S, Lyons SM, Kozakov D, Cifuentes D, Blower M, Kotton DN, Wilson AA, Muhlberger E, Emili A. 2020. Actionable Cytopathogenic Host Responses of Human Alveolar Type 2 Cells to SARS-CoV-2. Mol Cell 80:1104–1122 e9.

16. Stukalov A, Girault V, Grass V, Karayel O, Bergant V, Urban C, Haas DA, Huang Y, Oubraham L, Wang A, Hamad MS, Piras A, Hansen FM, Tanzer MC, Paron I, Zinzula L, Engleitner T, Reinecke M, Lavacca TM, Ehmann R, Wolfel R, Jores J, Kuster B, Protzer U, Rad R, Ziebuhr J, Thiel V, Scaturro P, Mann M, Pichlmair A. 2021. Multilevel proteomics reveals host perturbations by SARS-CoV-2 and SARS-CoV. Nature 594:246–252.

17. Borisova ME, Voigt A, Tollenaere MAX, Sahu SK, Juretschke T, Kreim N, Mailand N, Choudhary C, Bekker-Jensen S, Akutsu M, Wagner SA, Beli P. 2018. p38-MK2 signaling axis regulates RNA metabolism after UV-light-induced DNA damage. Nat Commun 9:1017.

18. Krischuns T, Gunl F, Henschel L, Binder M, Willemsen J, Schloer S, Rescher U, Gerlt V, Zimmer G, Nordhoff C, Ludwig S, Brunotte L. 2018. Phosphorylation of TRIM28 Enhances the Expression of IFN-beta and Proinflammatory Cytokines During HPAIV Infection of Human Lung Epithelial Cells. Front Immunol 9:2229.

19. Stokoe D, Engel K, Campbell DG, Cohen P, Gaestel M. 1992. Identification of MAPKAP kinase 2 as a major enzyme responsible for the phosphorylation of the small mammalian heat shock proteins. FEBS Lett 313:307–13.

20. Beltrao P, Albanese V, Kenner LR, Swaney DL, Burlingame A, Villen J, Lim WA, Fraser JS, Frydman J, Krogan NJ. 2012. Systematic functional prioritization of protein posttranslational modifications. Cell 150:413–25.

21. Hornbeck PV, Zhang B, Murray B, Kornhauser JM, Latham V, Skrzypek E. 2015. PhosphoSitePlus, 2014: mutations, PTMs and recalibrations. Nucleic Acids Res 43:D512-20.

22. Subramanian A, Tamayo P, Mootha VK, Mukherjee S, Ebert BL, Gillette MA, Paulovich A, Pomeroy SL, Golub TR, Lander ES, Mesirov JP. 2005. Gene set enrichment analysis: a knowledge-based approach for interpreting genome-wide expression profiles. Proc Natl Acad Sci U S A 102:15545–50.

23. Beardmore VA, Hinton HJ, Eftychi C, Apostolaki M, Armaka M, Darragh J, McIlrath J, Carr JM, Armit LJ, Clacher C, Malone L, Kollias G, Arthur JS. 2005. Generation and characterization of p38beta (MAPK11) gene-targeted mice. Mol Cell Biol 25:10454–64.

24. Raingeaud J, Whitmarsh AJ, Barrett T, Derijard B, Davis RJ. 1996. MKK3- and MKK6-regulated gene expression is mediated by the p38 mitogen-activated protein kinase signal transduction pathway. Mol Cell Biol 16:1247–55.

25. Schoggins JW, Rice CM. 2011. Interferon-stimulated genes and their antiviral effector functions. Curr Opin Virol 1:519–25.

26. Cuenda A, Rouse J, Doza YN, Meier R, Cohen P, Gallagher TF, Young PR, Lee JC. 1995. SB 203580 is a specific inhibitor of a MAP kinase homologue which is stimulated by cellular stresses and interleukin-1. FEBS Lett 364:229–33.

27. Kumar S, McDonnell PC, Gum RJ, Hand AT, Lee JC, Young PR. 1997. Novel homologues of CSBP/p38 MAP kinase: activation, substrate specificity and sensitivity to inhibition by pyridinyl imidazoles. Biochem Biophys Res Commun 235:533–8.

28. Johnson JL, Yaron TM, Huntsman EM, Kerelsky A, Song J, Regev A, Lin T-Y, Liberatore K, Cizin DM, Cohen BM, Vasan N, Ma Y, Krismer K, Robles JT, van de Kooij B, van Vlimmeren AE, Andrée-Busch N, Käufer N, Dorovkov MV, Ryazanov AG, Takagi Y, Kastenhuber ER, Goncalves MD, Elemento O, Taatjes DJ, Maucuer A, Yamashita A, Degterev A, Linding R, Blenis J, Hornbeck PV, Turk BE, Yaffe MB, Cantley LC. 2022. A global atlas of substrate specificities for the human serine/threonine kinome. bioRxiv doi:10.1101/2022.05.22.492882:2022.05.22.492882.

29. Hadfield J, Megill C, Bell SM, Huddleston J, Potter B, Callender C, Sagulenko P, Bedford T, Neher RA. 2018. Nextstrain: real-time tracking of pathogen evolution. Bioinformatics 34:4121–4123.

30. Jimenez-Guardeno JM, Nieto-Torres JL, DeDiego ML, Regla-Nava JA, Fernandez-Delgado R, Castano-Rodriguez C, Enjuanes L. 2014. The PDZ-binding motif of severe acute respiratory syndrome coronavirus envelope protein is a determinant of viral pathogenesis. PLoS Pathog 10:e1004320.

31. Kopecky-Bromberg SA, Martinez-Sobrido L, Palese P. 2006. 7a protein of severe acute respiratory syndrome coronavirus inhibits cellular protein synthesis and activates p38 mitogen-activated protein kinase. J Virol 80:785–93.

32. Padhan K, Minakshi R, Towheed MAB, Jameel S. 2008. Severe acute respiratory syndrome coronavirus 3a protein activates the mitochondrial death pathway through p38 MAP kinase activation. J Gen Virol 89:1960–1969.

33. Karki R, Sharma BR, Tuladhar S, Williams EP, Zalduondo L, Samir P, Zheng M, Sundaram B, Banoth B, Malireddi RKS, Schreiner P, Neale G, Vogel P, Webby R, Jonsson CB, Kanneganti TD. 2021. Synergism of TNF-alpha and IFN-gamma Triggers Inflammatory Cell Death, Tissue Damage, and Mortality in SARS-CoV-2 Infection and Cytokine Shock Syndromes. Cell 184:149-168 e17.

34. Pellegrina D, Bahcheli AT, Krassowski M, Reimand J. 2022. Human phospho-signaling networks of SARS-CoV-2 infection are rewired by population genetic variants. Mol Syst Biol 18:e10823.

35. Tovo PA, Garazzino S, Dapra V, Pruccoli G, Calvi C, Mignone F, Alliaudi C, Denina M, Scolfaro C, Zoppo M, Licciardi F, Ramenghi U, Galliano I, Bergallo M. 2021. COVID-19 in Children: Expressions of Type I/II/III Interferons, TRIM28, SETDB1, and Endogenous Retroviruses in Mild and Severe Cases. Int J Mol Sci 22.

36. Chang CK, Hou MH, Chang CF, Hsiao CD, Huang TH. 2014. The SARS coronavirus nucleocapsid protein--forms and functions. Antiviral Res 103:39–50.

37. Chen H, Gill A, Dove BK, Emmett SR, Kemp CF, Ritchie MA, Dee M, Hiscox JA. 2005. Mass spectroscopic characterization of the coronavirus infectious bronchitis virus nucleoprotein and elucidation of the role of phosphorylation in RNA binding by using surface plasmon resonance. J Virol 79:1164–79.

38. Wu CH, Chen PJ, Yeh SH. 2014. Nucleocapsid phosphorylation and RNA helicase DDX1 recruitment enables coronavirus transition from discontinuous to continuous transcription. Cell Host Microbe 16:462–72.

39. Savastano A, Ibanez de Opakua A, Rankovic M, Zweckstetter M. 2020. Nucleocapsid protein of SARS-CoV-2 phase separates into RNA-rich polymerase-containing condensates. Nat Commun 11:6041.

40. Wang J, Shi C, Xu Q, Yin H. 2021. SARS-CoV-2 nucleocapsid protein undergoes liquid-liquid phase separation into stress granules through its N-terminal intrinsically disordered region. Cell Discov 7:5.

41. Daniloski Z, Jordan TX, Wessels HH, Hoagland DA, Kasela S, Legut M, Maniatis S, Mimitou EP, Lu L, Geller E, Danziger O, Rosenberg BR, Phatnani H, Smibert P, Lappalainen T, tenOever BR, Sanjana NE. 2021. Identification of Required Host Factors for SARS-CoV-2 Infection in Human Cells. Cell 184:92–105 e16.

42. Seifert LL, Si C, Saha D, Sadic M, de Vries M, Ballentine S, Briley A, Wang G, Valero-Jimenez AM, Mohamed A, Schaefer U, Moulton HM, Garcia-Sastre A, Tripathi S, Rosenberg BR, Dittmann M. 2019. The ETS transcription factor ELF1 regulates a broadly antiviral program distinct from the type I interferon response. PLoS Pathog 15:e1007634.

43. Nilsson-Payant BE, Uhl S, Grimont A, Doane AS, Cohen P, Patel RS, Higgins CA, Acklin JA, Bram Y, Chandar V, Blanco-Melo D, Panis M, Lim JK, Elemento O, Schwartz RE, Rosenberg BR, Chandwani R, tenOever BR. 2021. The NF-kappaB Transcriptional Footprint Is Essential for SARS-CoV-2 Replication. J Virol 95:e0125721.

44. Torre D, Lachmann A, Ma’ayan A. 2018. BioJupies: Automated Generation of Interactive Notebooks for RNA-Seq Data Analysis in the Cloud. Cell Syst 7:556–561 e3.

45. Langmead B, Salzberg SL. 2012. Fast gapped-read alignment with Bowtie 2. Nat Methods 9:357–9.

46. Chen EY, Tan CM, Kou Y, Duan Q, Wang Z, Meirelles GV, Clark NR, Ma’ayan A. 2013. Enrichr: interactive and collaborative HTML5 gene list enrichment analysis tool. BMC Bioinformatics 14:128.

47. Chu DKW, Pan Y, Cheng SMS, Hui KPY, Krishnan P, Liu Y, Ng DYM, Wan CKC, Yang P, Wang Q, Peiris M, Poon LLM. 2020. Molecular Diagnosis of a Novel Coronavirus (2019-nCoV) Causing an Outbreak of Pneumonia. Clin Chem 66:549–555.

48. Bruderer R, Bernhardt OM, Gandhi T, Xuan Y, Sondermann J, Schmidt M, Gomez-Varela D, Reiter L. 2017. Optimization of Experimental Parameters in Data-Independent Mass Spectrometry Significantly Increases Depth and Reproducibility of Results. Mol Cell Proteomics 16:2296–2309.

49. Choi M, Chang CY, Clough T, Broudy D, Killeen T, MacLean B, Vitek O. 2014. MSstats: an R package for statistical analysis of quantitative mass spectrometry-based proteomic experiments. Bioinformatics 30:2524–6.

